# The monoaminergic system in a bivalve larva: temporal deployment and spatial organization

**DOI:** 10.64898/2026.08.17.745212

**Authors:** Beatrice Risso, Julie Blahuta, Lydia Besnardeau, Teresa Balbi, Remi Dumollard, Laura Canesi, Angelica Miglioli

## Abstract

Originating at the base of the bilaterian tree of life, the monoaminergic (MOA) system is a pivotal and evolutionarily conserved regulator of animal development and of responses to changing environmental conditions. Investigating the ontogeny of monoaminergic modulation in model systems such as marine bivalve molluscs is therefore particularly relevant, as their life cycle and developmental transitions are strongly influenced by environmental cues. Here, we characterized the spatio-temporal and tissue-specific expression of components of the MOA system during early larval development of the Mediterranean mussel *Mytilus galloprovincialis* using both time resolved transcriptomics and *in situ* Hybridization Chain Reaction (HCR). Our results identify serotonin and dopamine as the predominant and interconnected monoaminergic pathways deployed during early mussel development, with receptors, enzymes, and selective transporters broadly expressed across both neuronal and non-neuronal tissues. Notably, the expression of receptors preceding that of the corresponding biosynthetic enzymes indicates early, non-neuronal roles of monoaminergic signalling, supported by their localization in peripheral tissues such as ciliated epithelia. Altogether, These findings support the hypothesis that the MOA system acts as a pervasive and tightly regulated modulator of larval morphogenesis and could therefore play an evolutionary conserved role in mediating development and environmental plasticity in developing bilaterian organisms.

## Introduction

Originated at the base of the bilaterian tree of life, the monoaminergic system (MOA) is the ensemble of the cells and molecular elements required for the production, detection, transport, and degradation of monoamines (e.g., dopamine and serotonin), a class of low molecular weight biogenic amines derived from essential aromatic amino acids (1–5).

Monoamines are divided into four families according to their amino acid of origin (Fig. 1A): indolamines, imidazolamines, catecholamines and phenethylamines. Indolamines derive from L-tryptophan and include serotonin (5-HT) and melatonin; imidazolamines comprise histamine (HIS), derived from histidine; phenethylamines are trace amines synthesised from L-phenylalanine and include tyramine (TYM) and octopamine (OA); finally, catecholamines— including dopamine (DA), noradrenaline (NA) and adrenaline (A)—derive from L-tyrosine, a hydroxylated form of L-phenylalanine (6–10). Each monoamine defines a specific signalling circuit of the MOA system, comprising the neurons, target tissues, synthesis enzymes, vesicular and re-uptake transporters, G-protein coupled (GPCRs) and ion channel receptors, as well as the catabolic machinery necessary to tightly control availability and mediate neuronal and peripheral functions (Fig. 1B) (1, 4, 8–20).

**Figure 1.**
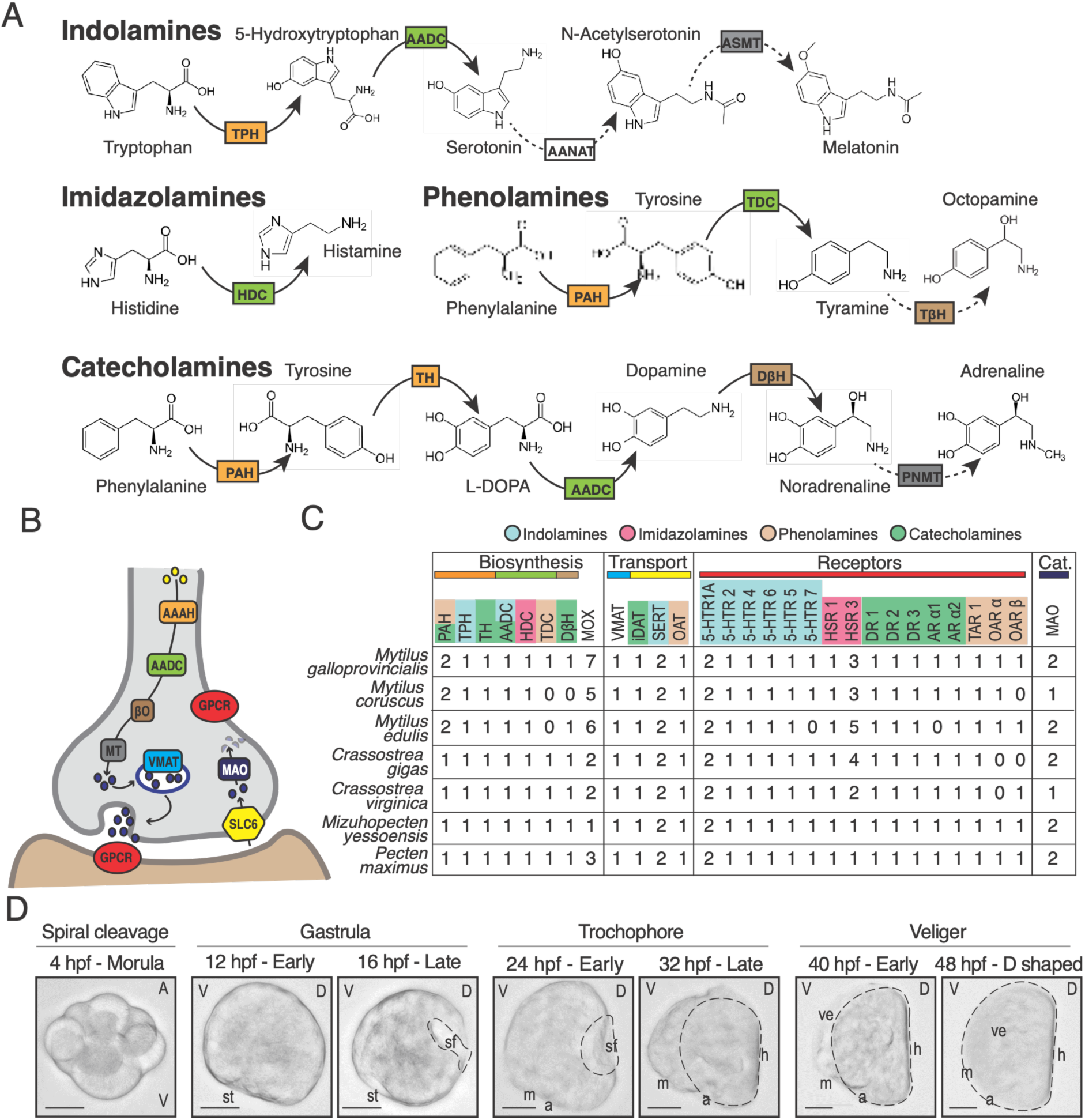
Overview of the molecular elements of the monoaminergic system of the Mediterranean mussel. A) Schematic representation of the synthesis pathway of the major monoamine neurotransmitters. Arrows indicate enzymatic reactions with the abbreviation of the enzyme overlayed. Dotted arrows indicate paths that are yet to be characterized or absent in bivalve molluscs. Abbreviations as in main text. Color code reflects the type of enzyme: orange – hydroxylases, green – decarboxylases, black – acetyltransferases, brown – β hydroxylases, grey - *N*-methyltransferases. B) Schematic representation of a monoaminergic neuron and monoamine target tissue displaying the pre-synaptic machinery regulating synthesis, storage, re-uptake, catabolism and reception of neurotransmitters. Colours as in panel A. C) Summary of orthology assessment analyses in *M. galloprovincialis* and other molluscs. Molecular elements are coloured by monoamine class, and abbreviations are as in the main text; colour bars reflect enzyme type as in A and B. Numbers indicate the number of proteins identified for each molecular element. D) Representative images of the embryonic and early larval development of the Mediterranean mussel. Staging as in Miglioli et al., 2025. In Morula: A - animal, V - vegetal. In all other panels: V – ventral, D – dorsal. St – stomodeum, m – mouth, a – anus, sf – shell field, h – hinge, ve – velum. Dotted line outlines the shell field. Scale bar: 20 µm.

In bilaterians studied so far, monoamines are generally produced by small populations of specialized (monoaminergic) neurons with extensive projections throughout the nervous system and peripheral organs, where they act as neurotransmitters, neurohormones, cytoskeletal and epigenetic regulators and form an integrative networks able to tune neuroendocrine signalling, cell cycle progression and gene expression in response to environmental inputs (1, 2, 21–24). Through this architecture, monoaminergic signalling can regulate/modulate locomotion, behaviour, stress responses, synaptic plasticity, as well as morphogenetic processes and developmental tempo (1–3, 7, 25, 26).

Comparative transcriptomic and developmental studies indicate that the architecture of the MOA system is highly homologous across Bilateria (1, 21, 22). In fact, despite the extensive lineage-specific diversification of neural architectures and neurotransmitter subtypes, bilaterians possess highly conserved biosynthetic and transporter machineries deployed by shared transcriptional program underpinning the development of monoaminergic neurons (1, 21, 22, 27). This deeply conserved molecular core operates early and across multiple aspects of bilaterian physiology by using both highly conserved and versatile regulatory networks (2, 23, 24) to mediate adaptive environmental plasticity during embryo-larval development (2, 23, 24).

However, comparative molecular analyses available almost exclusively focused on Ecdysozoans and Deuterostomes, overlooking Spiralia, the third main lineage of the bilaterian tree of life. Spiralians (e.g., molluscs, segmented worms) constitute an extremely diverse and species rich clade which include animals of environmental and economic interest such as marine bivalve molluscs (1, 21, 22, 27, 28). Characterized by planktonic embryos and intermediate larvae, their survival depends on the capacity to dynamically adapt to continuous fluctuations in pH, temperature and salinity of the water column (29, 30). Moreover, since they are devoid of a canonical endocrine system, the MOA system is a main actor in their neuroendocrine regulation, modulating, for example, ciliary beating, muscle contraction and relaxation, feeding, reproduction and shell formation (23, 24, 31, 32). Finally, both exposure to chemical contaminants and ongoing global changes such as ocean acidification have been shown to disrupt key components of monoaminergic signalling, affecting neuronal development and shell formation, as well as receptor and transporter expression, leading, to teratogenic effects and potential population declines (23, 33–40). While comparative neuroanatomical studies have provided important insights into the organization and evolution of bivalve nervous systems, consisting in a peripheral network progressively organized into ganglia, molecular information on the establishment of monoaminergic signalling remains fragmentary (41–46). 41-46). Recent work in M. galloprovincialis has begun to characterize the molecular organization of the serotonergic system, identifying tryptophan hydroxylase (TPH) and two serotonin reuptake transporters (SERT-like), describing their developmental expression, and showing their partial spatial colocalization in D-veliger larvae (36). However, these serotonergic components have not yet been placed within a comprehensive framework encompassing the different monoaminergic circuits, their developmental deployment, and their spatial organization across neuronal and peripheral tissues. Establishing such a framework in model bivalves would not only advance our understanding of the evolution of this regulatory system in Spiralia but also facilitate comparative and environmental studies investigating how monoaminergic pathways contribute to vulnerability and acclimation capacity of different bilaterian groups under global change scenarios.

To address this, here we combined time resolved transcriptomic and spatial transcript validation to provide the first characterization of the ontogeny of the MOA system of a bivalve mollusc, using the Mediterranean mussel *Mytilus galloprovincialis* as model system (47, 48). The results show that monoaminergic signalling is established early during embryogenesis across both neuronal and non-neuronal tissues, suggesting ancient functions that extend beyond canonical neurotransmission and include developmental regulation and environmental responsiveness. The coordinated, yet temporally distinct, deployment of serotonergic and dopaminergic systems, together with the early and widespread expression of monoamine receptors, suggests that the monoaminergic system constitutes an evolutionarily conserved interface linking morphogenesis, physiology, and adaptive plasticity across Bilateria.

## Methods

### Orthology assessment of monoaminergic elements in *M. galloprovincialis* genome

The molecular elements of the monoaminergic system (Fig. 1) of the Mediterranean mussel were retrieved from the Mg 10 genome assembly (49) by searching for proteins displaying the canonical domain architecture with Protein Basic Local Alignment Search Tool (BLASTP) on the Octopus Bioinformatic Server (https://octopus.obs-vlfr.fr/), using human (*Homo sapiens*) and fruit fly (*Drosophila melanogaster*) reference sequences as queries. Selected proteins by BLASTP were subjected to domain analysis using the NCBI Conserved Domain Database (CDD) (50, 51). Only sequences containing the expected functional domains were retained for further analysis. Orthology was assessed by Maximum Likelihood (ML) with MEGA (MEGA-X 10.2.6) with 1000 bootstraps prior multiple sequence alignment with ClustalW (BioEdit version 7.0.2) (52, 53). Substitution model selection and rate estimation were performed for each dataset of protein sequences using MEGA-X internal tools. When ML analysis did not yield interpretable results, orthology was determined based on presence/absence of amino acids involved in the catalytic site or in ligand binding through FASTA sequence alignment using ClustalW (BioEdit version 7.0.2) (53). To strengthen orthology assignments, the same analyses were performed on homologous sequences from other bivalve species with public genome repositories in NCBI following the selection criteria outlined in Canesi et al. (2022): *Mytilus edulis*, *Mytilus coruscus*, *Crassostrea gigas*, *Crassostrea virginica*, *Mizuhopecten yessoensis* and *Pecten maximus*.

### Transcriptomic analysis of the elements of the monoaminergic system in *M. galloprovincialis* embryo-larval development

Developmental expression of monoaminergic genes was analysed using the reference developmental transcriptome of the Mediterranean mussel (47), selecting libraries spanning embryonic and larval development (0–48 hpf) (Fig. 1D). Transcript abundance values were expressed as Transcripts Per Million (TPM) and Z-scored to visualise relative temporal expression dynamics across developmental stages. Genes of interest were identified based on MGAL gene identifiers and assigned to monoaminergic pathways according to functional annotation. Temporal expression profiles were visualised as heatmaps using the R package pheatmap (54). Genes were grouped according to monoaminergic pathway, and developmental stages were annotated as previously described (47). To summarise pathway-level developmental trends, mean Z-scored expression values were calculated for each monoaminergic pathway across developmental stages and fitted using locally estimated scatterplot smoothing (LOESS) in ggplot2 (54). Shaded ribbons represent the uncertainty associated with the fitted LOESS trend, while stars indicate the developmental stage at which each pathway reached its maximum mean expression.

### Spawning and fertilization

Sexually mature specimens of *M. galloprovincialis* were sampled from a natural population in the bay of Villefranche-sur-Mer (43.682°N, 7.319°E, France) during the spawning season (January-March 2023). Mussels were acclimatized in flow-through vessels containing filtered natural seawater (0.2 μm Millipore filter), pH 8.0-8.2, 38 ppt salinity, 15°C (Millipore filtered sea water, MFSW) by the Centre de Ressources Biologiques Marines (CRBM) at the Institut de la Mer de Villefranche (IMEV). Gametes were obtained by heat-shock as previously described (55). Egg quality and sperm motility were checked by microscopy observations; fertilization and assessment of fertilization success were performed as previously reported (48, 55). After 30 min, fertilized eggs were transferred to culture vessels at a density of 200 larvae/mL (filtered natural seawater, pH 8.0-8.2, 16°C).

### Pharmacology and larval sampling

Following standard embryotoxicity assay guidelines (56), 30 mins post-feralization embryos were exposed to agonists and antagonists of the serotonergic and dopaminergic systems and the effects were evaluated on the formation of D-veliger at 48h. Experiments were conducted in 24 well plates and 50 mL flasks as previously described (33, 34, 55). Test substances included: serotonin (5-HT, stock solution: 100 mM in DMSO, Sigma-Aldrich,153-98-0), dopamine (DA, stock solution: 100 mM in DMSO, Sigma-Aldrich, 62-31-7), methiothepin, a non-selective inhibitor of serotonin receptors, (MT, stock solution 100 mM in DMSO, Sigma-Aldrich 74611-28-2) and SCH 23390, a selective inhibitor of dopamine receptor 1 (DR1), (SCH, stock solution 50 mM in DMSO, Sigma-Aldrich, 125941-87-9) (34, 35). All chemicals were diluted in MFSW (Millipore Filtered Seawater) to obtain the final desired concentrations. DMSO (Dimethyl Sulfoxide) at the highest concentration (0.01%) was added to controls. A minimum of four parental pairs were analysed for each condition. Samples for morphological analyses and *in situ* hybridization chain reaction (HCR) were collected at 16, 28 and 48 hpf using a 0.2 μm filter and fixed in 4% paraformaldehyde (PFA) in 1× Phosphate Buffered Saline (PBS, 137 mM NaCl, 2.7 mM KCl, 10 mM Na_2_HPO_4_, 1.8 mM KH_2_PO_4_, pH 7.4) solution overnight at 4°C. Samples were then washed 3 times for 15 minutes in 1× PBS and stored in 100% methanol at −20 °C (47).

### Morphological analysis of larvae

Morphological analyses were performed at 48 hpf (D-Veliger 2): according to the ASTM (2012) larvae were considered normal when they were D-shaped, with a straight hinge and an absence of protruding mantle. Larvae were considered malformed when protruding mantle or/and hinge malformations were present and delayed/arrested when they were at earlier stages (pre-veliger or trochophore) (34, 48). Data on larval phenotypes are reported as the percentage of normal over malformed D-Veliger larvae, evaluated in at least 50 larvae per each parental pair and experimental condition, each analysed independently. Brightfield images were acquired with 20× objective using the inverted microscope Axio Observer 7. Statistical analysis of the observed effects was performed using Kruskal-Wallis test and Dunn test for *post-hoc* comparison with Bonferroni-adjusted p-values to show differences between controls and treatments in terms of percentage of normal D-Larvae (ND).

### *In situ* Hybridization Chain Reaction (HCR)

Probe sets for *in situ* HCR (Tab. S1) were designed using a user-friendly Python interface (https://github.com/rwnull/insitu_probe_generator) and synthesized by OligoPools (Twist Bioscience) as previously described (47). Amplifiers were purchased from Molecular Instruments (https://www.molecularinstruments.com). Synthesis enzymes and degradation enzymes genes were marked with amplifier in yellow (B2, ex: 561 nm, em: 572 nm, laser: 552 nm), receptors with amplifier in red (B1, ex: 650 nm, em: 671 nm, laser: 638 nm) and transporters with amplifier in green (B3, ex: 518 nm, em: 543 nm, laser: 488 nm). To establish the localisation of genes of interest (Tab. S1), they were co-labelled with validated tissue-specific marker genes for secretory neurons (*secretogranin V/7b2*), ciliated epithelium (*Tektin/Tek*), muscle system (*Myosin heavy chain/Mhc*) (Tab. S2) (47, 57).The protocol was carried out as previously described (47).

### Image acquisition and analysis

Specimens were mounted in the antifading agent CitiFluor AF1 (Agar Scientific) and imaged with a SP8 confocal microscope (Leica Microsystem) with LasX software with the following settings: three-dimensional acquisition mode (xyz), bidirectional scanning mode, 1024 × 1024 image format, scanning speed of 600 Hz, pixel size of 200 nm × 200 nm, z-step size of 0.5 µm for single HCR or 0.8 µm for double/triple HCR, line accumulation and frame average set to 3 and sequential mode to avoid spillover between the different channels. Pictures were taken with 40× objective and a minimum of five larvae were analysed for each gene/gene combination. Images were analysed with Fiji (58). “Math::Subtract” and “Process::Subtract Background” were applied uniformly across all samples to remove background noise and smooth signal. Selection of planes were used to obtain MAX and 3D projections with default settings to annotate tissue-specific expression and assess co-expression regions. Specimen orientation was adjusted with “Image::Transform”.

## Results

### Characterization of components of the monoaminergic system in *M. galloprovincialis*

Orthology assessment of the molecular elements of the monoaminergic system in *M. galloprovincialis* genome revealed the presence of up to six monoaminergic circuits, including serotonin, dopamine, noradrenaline, tyramine, octopamine and histamine pathways (Fig. 1C). These systems comprise the expected components required for synthesis, transport, reception and degradation, consistent with the conserved architecture of monoaminergic signalling in protostomes (1).

#### a) Synthesis enzymes

Aromatic amino acid hydroxylases (AAAH) are the first rate limiting enzymes for the synthesis of all Indolamines, Catecholamines and Phenolamines (Fig. 1A, B), and include, respectively: tryptophan hydroxylase (TPH), tyrosine hydroxylase (TH) and phenylalanine hydroxylase (PAH) (1). As previously reported, four AAAH sequences are present in the *M. galloprovincialis genome*, corresponding to one TPH, one TH and two PAHs (36). The same complement was identified in the other Mytilidae analysed, whereas the second PAH was not detected in the other bivalve species examined (36). These previously characterized components were incorporated here into the broader reconstruction of the monoaminergic repertoire (Fig. 1C).

Next, aromatic L-amino acid decarboxylases (AADC) were investigated. Decarboxylases catalyse the removal of a carboxyl group and include: aromatic L-amino acid decarboxylase (AADC), the enzyme converting 5-hydroxytryptophan in serotonin and L-DOPA in dopamine; histidine decarboxylase (HDC), which converts histidine to histamine, and tyrosine decarboxylase (TDC), which is responsible for the conversion of tyrosine to tyramine (Fig. 1A, B) (1, 59–62). Orthology assessment identified one representative of each AADC in all investigated species (Tab. S3, Fig. S1, Fig. 1C), with relatively high support (bootstrap values – BP > 70), except TDC, which displayed slightly lower branch support (BP=59). Altogether, the AAAHs and ADDCs identified confirm that bivalves can synthesise serotonin, dopamine, tyramine and histamine (Fig. 1A, B, C).

Then, sequences displaying copper-containing hydroxylase domains were searched to identify: dopamine-β-hydroxylase (DβH), that catalyse the hydroxylation of dopamine to noradrenaline (or norepinephrine), tyramine-β-hydroxylase, which converts tyramine in octopamine, and monooxygenase X (DBH-like monooxygenase protein 1 – MOXD1) whose function is still unknown but speculatively associated to the catecholaminergic circuit (1, 13, 15).

Orthology assessment revealed the presence of one DBH in bivalve molluscs (BP = 75), except for *M. coruscus*, in which none was detected probably due to genome uncompletedness, the presence of several MOXD1 (here called DBH-like, BP = 99) and, in line with previous reports, no TBH, which is specific to Ecdysozoans (Tab. S4, Fig. S2, Fig. 1C) (1, 32).

Finally, no bivalve sequences were found for phenylethanolamine-N-methyltransferase (PNMT), the enzyme responsible for the synthesis of adrenaline from noradrenaline, serotonin N-acetyltransferase (AANAT) and Acetylserotonin O-Methyltransferase (ASMT), responsible for the conversion of serotonin into melatonin, confirming previous observations (1, 32).

#### b) Degradation enzymes

Monoamines are catabolised by monoamine oxidases (MAO) A and B. As orthology assessment did not yield clear results in either *Mytilus* or the other bivalve molluscs (Tab. S5, Fig. S3), multiple sequence alignment was carried out to search for the presence/absence of the amino acids responsible for monoamine catabolism: phenylalanine 208 (F208) for MAO-A, and isoleucine 199 (I199) for MAO-B (63, 64). To perform such analysis, murine sequences were also included. The results led to the classification of *M. galloprovincialis* MAOs as MAO-A like (Fig. S4A) and MAO-B (Fig. S4B). The first one shows a similar amino acid (tyrosine) aligning with human and murine MAO-A, whereas the second presents an isoleucine like human and murine MAO-B (Fig. 1B, C, Tab. S4). In the other species of bivalve molluscs analysed, at least one MAO or MAO-like sequence was found (Tab. S6).

#### c) Transporters

Membrane transporters of the Solute Carrier (SLC) family regulate monoamine availability through reuptake (SLC6) and vesicular storage and release (SLC18) (Fig. 1B). SLC6 neurotransmitter/sodium symporters mediate the reuptake of serotonin (SERT), dopamine (DAT), noradrenaline (NET), tyramine and octopamine (OAT) at neuronal and peripheral interfaces (1, 17, 18). The SLC6 repertoire of *M. galloprovincialis* and other bivalves has been previously characterized, revealing two SERT-like transporters, an octopamine transporter (OAT) and an invertebrate dopamine transporter (iDAT), while canonical NET and histamine transporters were not identified (36). These previously characterized transporters were incorporated here into the reconstruction of the corresponding monoaminergic circuits (Fig. 1B, C).

The second main transporters controlling monoamine storage and release are SLC18-type, which operate the predominantly pre-synaptic vesicular transport (VMAT) of all major monoamines. Orthology assessment for SLC18s was performed using the sequences reported in Tab. S7. All protostomes analysed displayed a single VMAT (Fig, 1B, C; Fig. S5) as previously reported for invertebrates at large (17, 65).

#### d) Receptors

Monoamines exert their action by binding to G-protein-coupled receptors (GPCRs) in the pre- and post-synaptic membranes. Regarding serotonin, at least three different 5-HTRs have been so far identified in bivalve molluscs: 5-HTR1A1, 5-HTR1A2 and 5-HTR7 (23). Orthology assessment revealed the presence of six serotonin receptor types in bivalves: 1A, 2, 4, 5, 6 and 7. 5-HTR7 was not found for *M. edulis*, probably due to low genome resolution (Fig. 1C, Tab. S8, Fig. S6). In line with previous studies, no 5HTR3-type receptors were found (1). Regarding dopamine, a recent study defined three receptor types in invertebrates: type 1 (DR1), type 2 (DR2, or invertebrate dopamine receptor – INDR), and type 3 (DR3-LIKE) (66) (Tab. S9, Fig. S7). Orthology assessment confirmed these results in *M. galloprovincialis* and the other bivalve species, with the exception of *M. coruscus*, for which DR1 could not be identified, probably due to low genome resolution (Fig. 1C, Fig. S7).

For adrenergic, tyramine and octopamine receptors, reference sequences from *Platynereis dumerilii* (annelid) were used as additional query (Tab. S10) (27). Analyses revealed that *M. galloprovincialis* and other bivalves present one or two adrenergic α receptors (ARα) and octopamine receptors (OARs) depending on the species, and one tyramine receptor (TARs) (Fig. 1C, Fig. S8). In *M. galloprovincialis*, two ARs, two OARs and one TAR were found (Fig, 1C, Fig. S8). As expected, the pacific oyster did not show any OAR and no ARβ was found outside the vertebrate species analysed (27, 66).

For histamine receptor (HSR), sequences from the acorn worm *Saccoglossus kowalevskii* (hemichordate) were used as reference (67). All bivalves displayed one HSR1 and a variable number of HSR3 (three in *M. galloprovincialis*). No HSR2 was found in any of the bivalves analysed (Tab. S11, Fig. 1C, Fig. S9).

### Developmental expression of monoaminergic genes in *M. galloprovincialis*

To investigate the temporal deployment of monoaminergic pathways during development, expression dynamics of the identified monoaminergic genes were examined across the first 48 hours of development, from fertilization to the D-veliger stage, using the reference developmental transcriptome of M. galloprovincialis (47). Developmental expression of *tph*, *sert1-like* and *sert2-like* had been previously analysed individually (36); here, these genes were integrated with the broader monoaminergic repertoire to reconstruct the temporal deployment of the different signalling circuits. (Fig. 2A, B). Expression profiles revealed that the molecular components of each monoaminergic pathway are assembled progressively throughout development rather than being activated simultaneously (Fig. 2A). Several receptors and transporters were already expressed during embryogenesis, whereas biosynthetic enzymes and additional signalling components frequently appeared at later developmental stages. For example, histamine (*hsr1-like*), tyramine (*tar1*), serotonin (5*htr1a1-2*) and dopamine receptors (*dr3-like*) were detected prior to the maximal expression of their corresponding biosynthetic pathways, suggesting that monoaminergic signalling may initially operate through maternally supplied monoamines, environmental ligands, or non-neuronal signalling mechanisms. Similarly, the temporal separation between receptors, transporters andsynthetic enzymes indicates that individual monoaminergic circuits are established through multiple developmental waves or built gradually through several distinct stages of development, rather than a single coordinated activation event.

**Figure 2.**
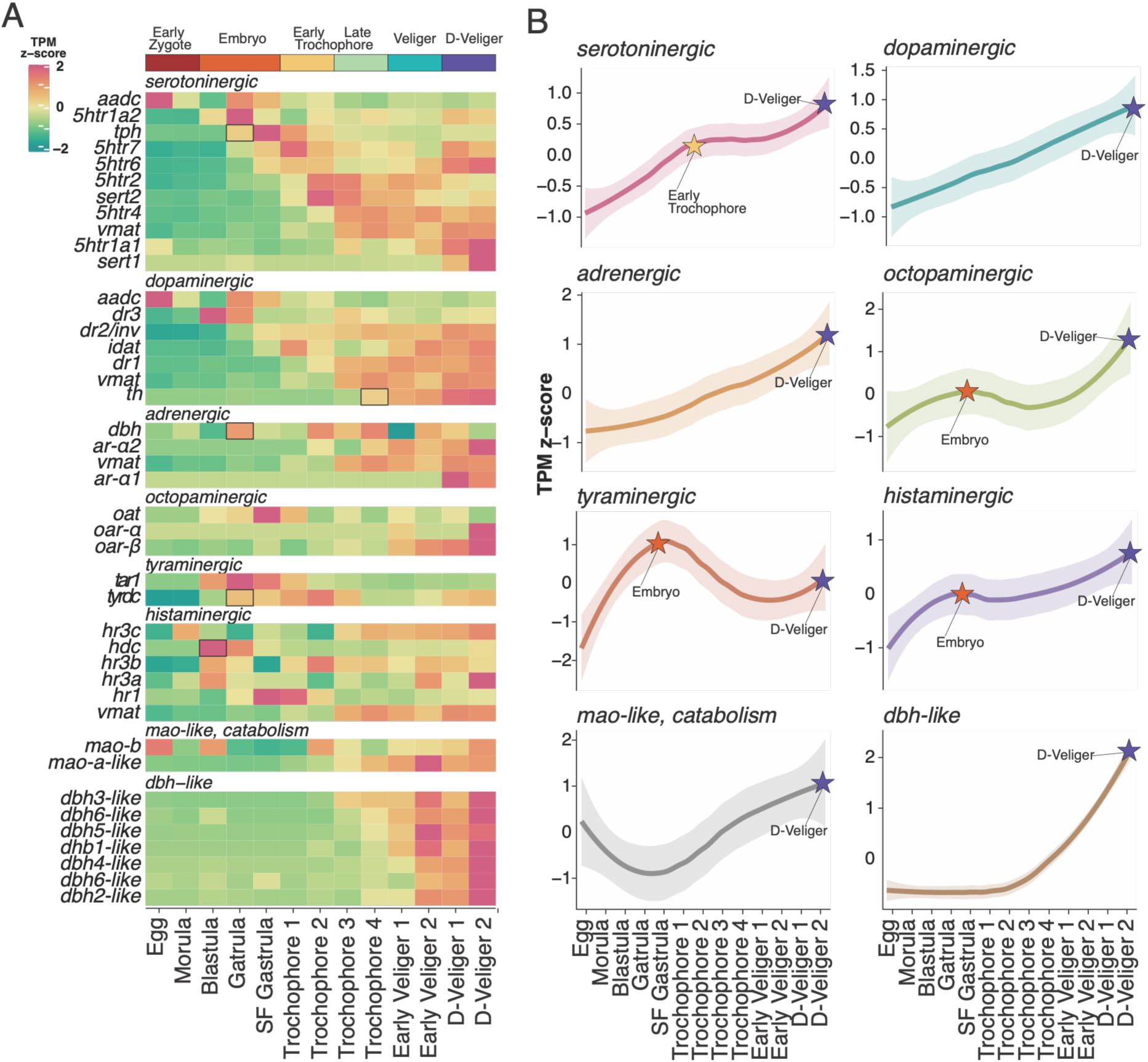
Developmental expression of monoaminergic genes in M. galloprovincialis. A) Clustered heatmaps of the expression of the elements of each monoaminergic circuit. Black outlines indicate first detectable expression of the rate limiting enzyme for each monoaminergic circuit. B) Mean expression of each monoaminergic circuit across development, including the catabolic machinery (*mao*) and the presumptive *dβh-like/mox*. Stars indicate the peaks of expression of the circuit.

To summarise the overall developmental deployment of each pathway, mean expression trajectories were reconstructed at the circuit level (Fig. 2B). Distinct temporal patterns emerged. The tyraminergic circuit reach maximal expression during embryonic development, while histaminergic and octopaminergic ones reached a first increase in expression during embryogenesis, followed by a subsequent peak by the D-Veliger stage, indicating an early and late developmental role for these pathways. Serotonergic signalling peaked during early trochophore stages, whereas dopaminergic, adrenergic, monoamine catabolic (*mao-like*) and *dopamine β-hydroxylase-like* pathways displayed maximal expression during veliger and D-veliger development (Fig. 2B). These results indicate a progressive shift in monoaminergic signalling across ontogeny, with early deployment of tyraminergic, histaminergic and octopaminergic pathways followed by serotonergic activation and the subsequent enrichment of catecholaminergic circuits during larval maturation. Together, the gene-level and pathway-level analyses reveal that monoaminergic pathways are assembled progressively throughout development while displaying distinct periods of maximal deployment, supporting an increasing complexity of monoaminergic regulation as larval tissues differentiate. Because of gene expression levels and circuit completeness, only 5-HT and DA systems were furtherly investigated by *in situ* HCR.

### The neurosecretory system of *M. galloprovincialis* larvae

The larval neurosecretory system of the Mediterranean mussel was characterized from Late Gastrulae to D-Veligers by double *in situ* HCR with Secretogranin V (*7b2*), a neuroendocrine protein detected in the secretory neurons of both deuterostomes and protostomes (68, 69) (Fig. 3A and B). To establish the topology of neural tissues, *7b2* was first multiplexed with the muscle marker *myosin heavy chain (mhc)* and the ciliated epithelium marker *tektin-3* (*tek*) (47).

**Figure 3.**
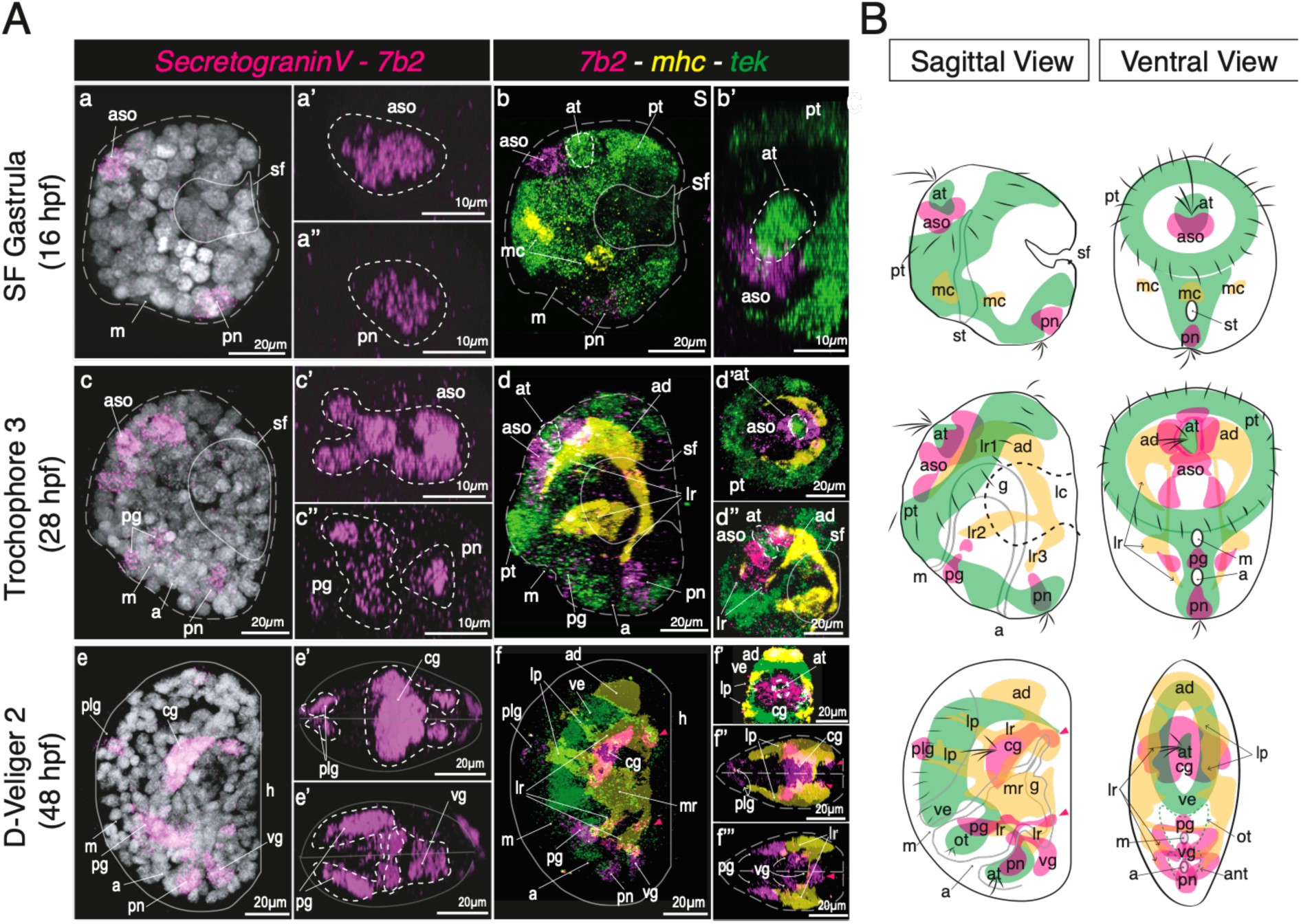
The larval neurosecretory system of the Mediterranean mussel. A) *In situ* HCR images of *7b2* (magenta) expression and co-expression with the muscle marker *mhc* (yellow) and ciliated epithelium marker *tek* (green) in *M. galloprovincialis* larvae. a, c, e) Sagittal views of 7b2 expression; a’-‘’, c’-‘’,e’-‘’) anterior views of the, respectively, anterior and posterior neurosecretory ganglions; b, d, f) Sagittal views of the co-expression of *7b2, mhc*, and *tek*; b’) Anterior view of the apical sensory organ with *7b2* and *tek;* d’-‘’) Anterior and lateral views of the apical sensory organ with *7b2, tek* and *mhc*; f’) Ventral view of the cerebral ganglion with *7b2*, *mhc* and *tek*; f’’-‘’’). Aso: apical sensory organ; at: apical tuft; sf: shellfield; vg: visceral ganglion; m: mouth; pt: prototroch; mc: muscle cells; pg: pedal ganglion; pn: posterior neurons; ad: anterior adductor muscle; lc: larval muscle chord; lr: larval retractor muscles; lp: larval protractor muscles; h: hinge; cg: cerebral ganglion; plg: pleural ganglion; a: anus; m: mouth; st: stomodeum; g: gut; ot: oral tuft; an: anal tuft. Dotted grey lines outline the larval body, dotted white lines outline the apical tuft, continuous lines outline the growing shell and shellfield. Scalebars as indicated. B) Schematic representation of *M. galloprovincialis* neurosecretory system (sagittal and ventral views) with respect to muscle system (yellow) and ciliated epithelium (green).

At the end of gastrulation (at 16 hpf), two populations of neurosecretory (*ns*) cells were detected: one anterior cluster in the presumptive apical sensory organ (*aso*) proximal to the apical tuft (*at*) (Fig. 3 b-b’), and a second cluster at the posterior end of the embryo (called *pn* for posterior neurons in Fig. 3 a-a’’). At the Mid Trochophore stage (28 hpf), the *ns* cluster of the apical sensory organ has expanded and surrounds the apical tuft on its ventral side and is enclosed by the most anterior pair of larval retractor muscles (lr) stemming from the anterior adductor muscle (Fig. 3 c, c’, d, d’, d’’). The posterior ns cells (*pn*) are opposed to the third pair of larval retractors (*lr*) (Fig. 3 c, c’’, d). A third population appears in the medial ventral ectoderm in the presumptive pedal ganglion (*pg*) (Fig. 3 c, d). By the D-Veliger stage (48 hpf), the anatomy of the larva is significantly changed, as the muscle system retracts the larval prototroch within the D-shaped shell, causing the anterior ventral ectoderm, including the aso, to fold (velum, ve) inside the shell. The ns cells located in the aso, now called cerebral ganglion (*cg*), still surround the apical tuft and are anchored to the anterior adductor muscle that forms a muscle ring (Fig. 3 e, e’, f, f’, f’’). The posterior neurons and pedal ganglion are now connected through the visceral ganglion (*vg*) around the mouth (*m*) and are anchored to the muscle chord connecting the two sides of the posterior muscle system (Fig. 3 e, e’’, f’’’). Finally, a new population of bilateral neurosecretory cells appear at the ventral end of the larval protractor muscles in the presumptive pleural ganglion (*plg*, Fig. 3 e, e’, f, f’, f’’).

### Expression of serotoninergic pre-synaptic machinery

Serotoninergic pre-synaptic machinery, *sensu stricto*, comprises the rate limiting enzyme for 5-HT production *tph*, the decarboxylase *aadc* for serotonin production and the transporters *vmat* (for vesicular storage) and *sert1-2* for re-uptake. These elements were studied by multiplexed *in situ* HCR with *7b2* to determine their precise neural location (Fig 4). The results show that ns 5-HT producing cells (expressing *tph* and *aadc*) are detected already in late gastrula in the aso. *aadc* is also detected more broadly, although with a weaker signal, in the ventral ectoderm of late gastrulae and gut region of trochophores and D-Veligers. In contrast, expression of *tph* remains restricted to ns cells of the *aso/cg* on the ventral side of the apical tuft in both trochophore and D-Veligers. The vesicular transporter *vmat* co-localizes with *tph* in ns cells throughout larval development and displays additional expression in the pedal ganglion (*pg*). *Sert2* is expressed in the medial section of the tph cell cluster in all stages analysed and, additionally, in ciliated epithelia. *Sert1* is specifically detected in the lateral *tph* ns cells of the cerebral ganglion of D-Veliger larvae, displaying no overlap with *sert2*. Overall, the imaging data suggest that mussel larvae can already, virtually, produce and store 5-HT at late gastrulation, while re-uptake starts during the trochophore stage.

**Figure 4.**
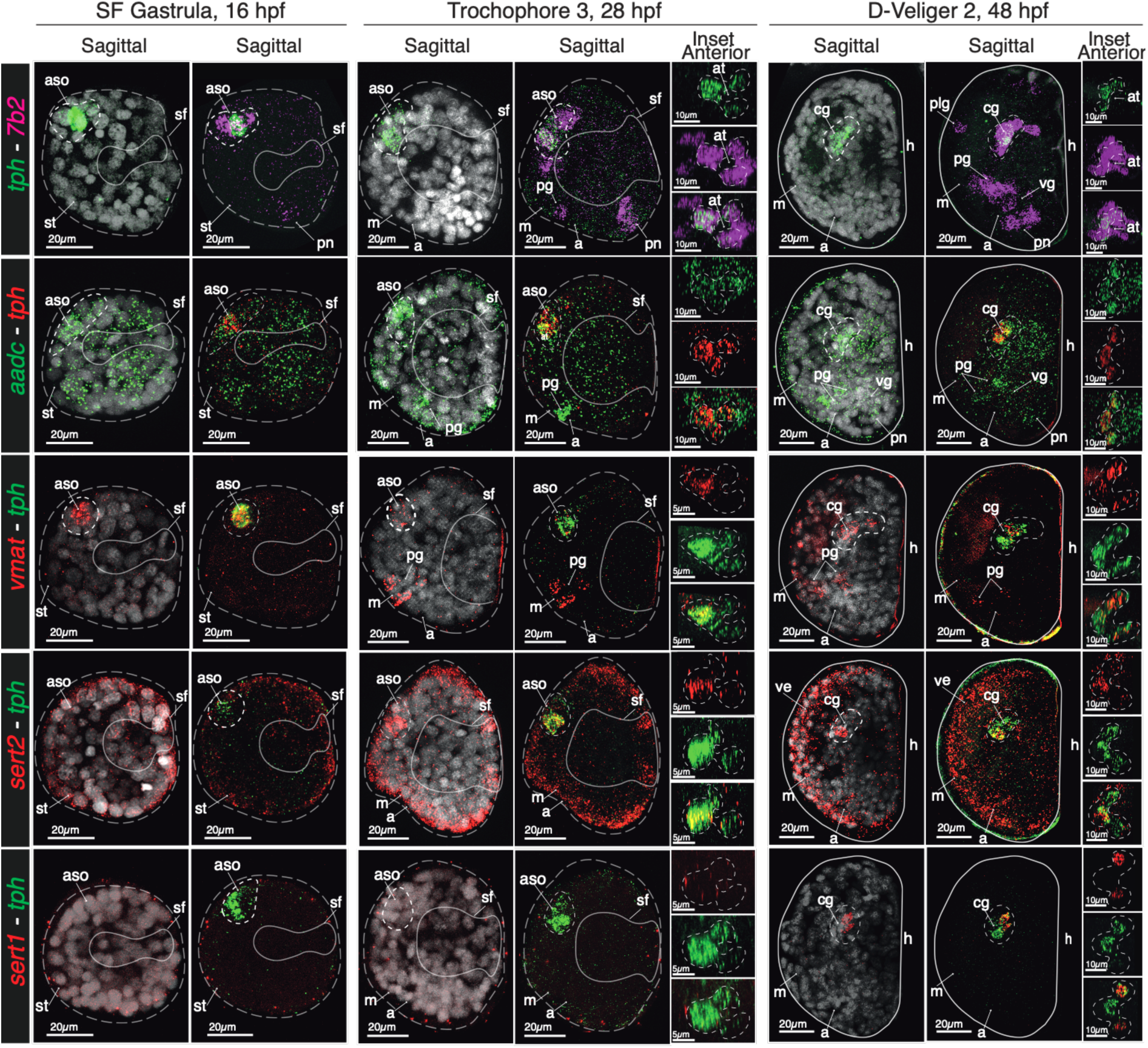
The serotoninergic pre-synaptic machinery in *M. galloprovincialis* larvae. Expression of pre-synaptic serorotoninergic markers in late gastrulas, mid trochophores and D-Veligers. Genes and colour codes as indicated in lateral bars: *tph – tryptophan hydroxylase, 7b2 – secretogranin V, aadc – aromatic aminoacid decarboxylase; vmat - vescicular amine transporter; sert1/2 – serotonin re-uptake transporter 1* and *2*. Insets display the anterior view of the apical sensory organ/cerebral ganglion with the single and multiplexed genes channels. aso: apical sensory organ; at: apical tuft; m: mouth; a: anus; st: stomodeum; sf: shellfield; cg: cerebral ganglion; h: hinge; ve: velum; pg: pedal ganglion; plg: pleural ganglion; vg: visceral ganglion; pn: posterior neurons. Dotted grey lines outline the larval body, continuous grey lines outline the growing shell, dotted white lines outline the area of the apical sensory organ. Scalebars as indicated.

### Expression of dopaminergic pre-synaptic machinery

The dopaminergic pre-synaptic machinery comprises the rate limiting enzyme for DA production *th*, the decarboxylase *aadc*, and the vesicular and re-uptake transporters *vmat* and *idat*, respectively. Using the same approach and rationale as for 5-HT, the elements were studied by multiplexed *in situ* HCR with *7b2* and *th* to gain insight into the onset of DA neurotransmission (Fig 5A). The results show that ns DA producing cells are detected starting from the trochophore stage in the pedal ganglion and *th* expression remains restricted to the *pg* in D-Veligers. The co-expression of *th* with *aadc* is suggestive of active dopamine production during these stages. Both *vmat* and *idat* also display clear expression in the most ventral *th* positive clusters in both larval stages (Fig. 5A). Interestingly, *idat* showed expression in non-neuronal regions that were recognized as part of the developing adductor anterior muscle and larval retractor muscles by colocalization with *mhc* (Fig. 5B). In addition, a set of DBH-like transcripts identified through orthology analysis was examined (data not shown). The paralogs displayed overlapping spatial expression patterns in the larval gut, consistent with the one previously described (34).

**Figure 5.**
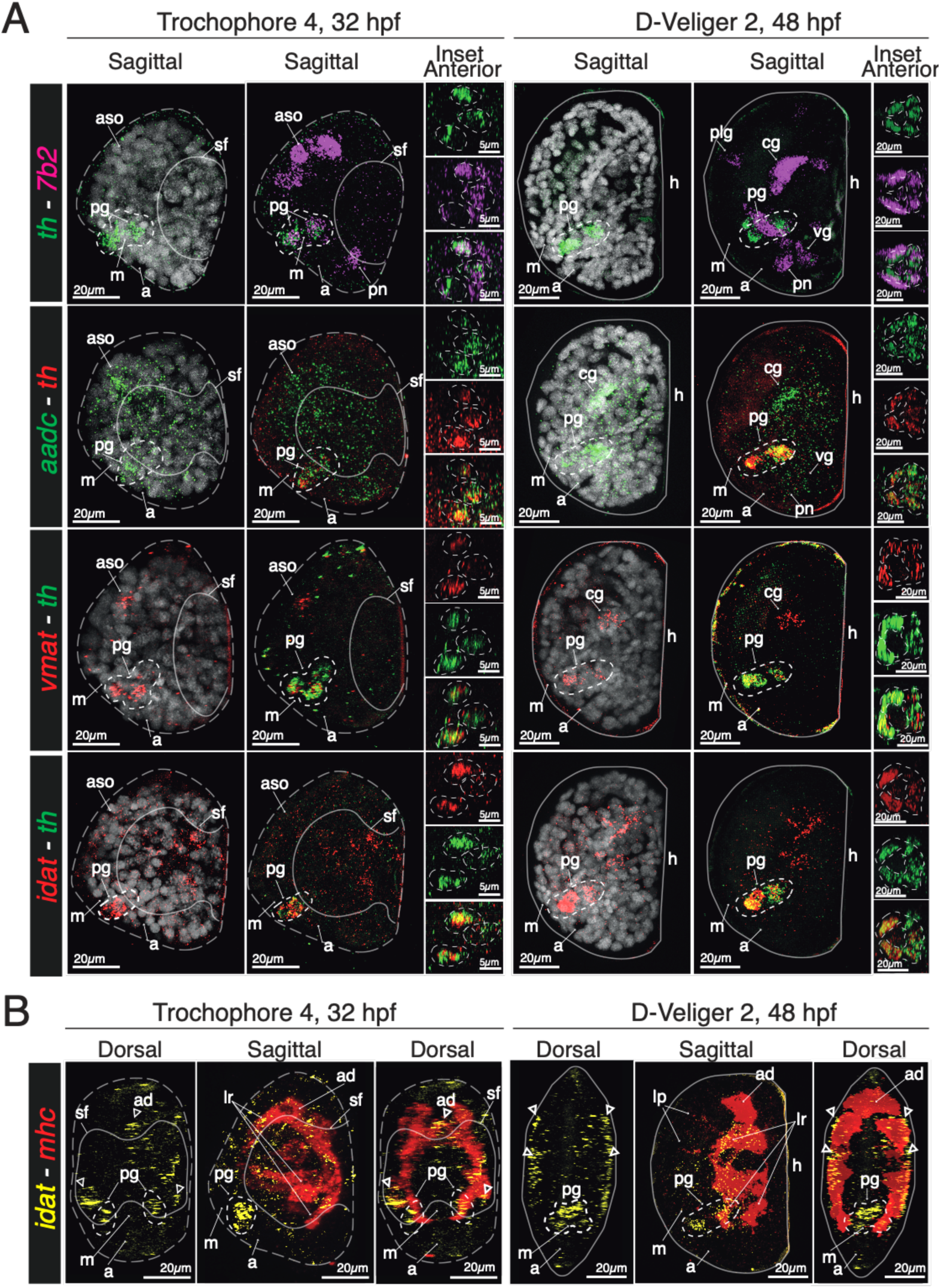
The dopaminergic pre-synaptic machinery in *M. galloprovincialis* larvae. A) Expression of pre-synaptic dopaminergic markers in Trochophore 4 and D-Veligers. Genes and colour codes as indicated in lateral bars: *th – tyrosine hydroxylase, 7b2 – secretogranin V, aadc – aromatic aminoacid decarboxylase; vmat - vescicular amine transporter; idat – invertebrate dopamine re-uptake transporter*. Insets display the anterior view of the pedal ganglion with the single and multiplexed genes channels. B) Expression of *idat* and *mhc* in Trochophore 4 and D-Veligers. aso: apical sensory organ; m: mouth; a: anus; sf: shellfield; cg: cerebral ganglion; h: hinge; ve: velum; pg: pedal ganglion; plg: pleural ganglion; vg: visceral ganglion; ad: adductor anterior muscle; lr: larval retractor muscles; pl: larval protractor muscles. Dotted grey lines outline the larval body, continuous grey lines outline the growing shell, dotted white lines outline the area of the pedal ganglion. White arrowheads indicate the non-neuronal expression of *idat* in the muscle system. Scalebars as indicated.

### Expression of serotonin receptors

Next, we studied the developmental expression pattern of 5-HT receptors (Fig 6, 7). *5-htr1a2* displayed expression in the *tph* positive cells of the *aso* and *cg* throughout the stages analysed, with expression in the pedal ganglion appearing in trochophores (Fig. 6A). *5-htr1a1* showed the same localization but, consistent with RNA-seq data, it was detectable only in D-Veliger larvae (Fig. 6B). Conversely, *5-htr4* localized in the regions of the posterior neurons and visceral ganglion, with weak expression in the pedal ganglion (Fig. 6C). The non-neuronal expression of *5-htr4* was further investigated by colocalization with the muscle marker *pmyo* (paramyosin), revealing some level of overlap between the receptor and the anterior portion of the larval muscle system in trochophores and D-Veligers (Fig. 6D). *5htr6* was consistently expressed in the posterior neurons of the visceral ganglion throughout larval development, with some weak localization in the *cg* and *pg* of D-Veliger larvae (Fig 7A). Conversely, *5htr7* displayed no expression in ns cells but was instead expressed in the ciliated epithelia of developing larvae including prototroch and velum (Fig 7B).

**Figure 6.**
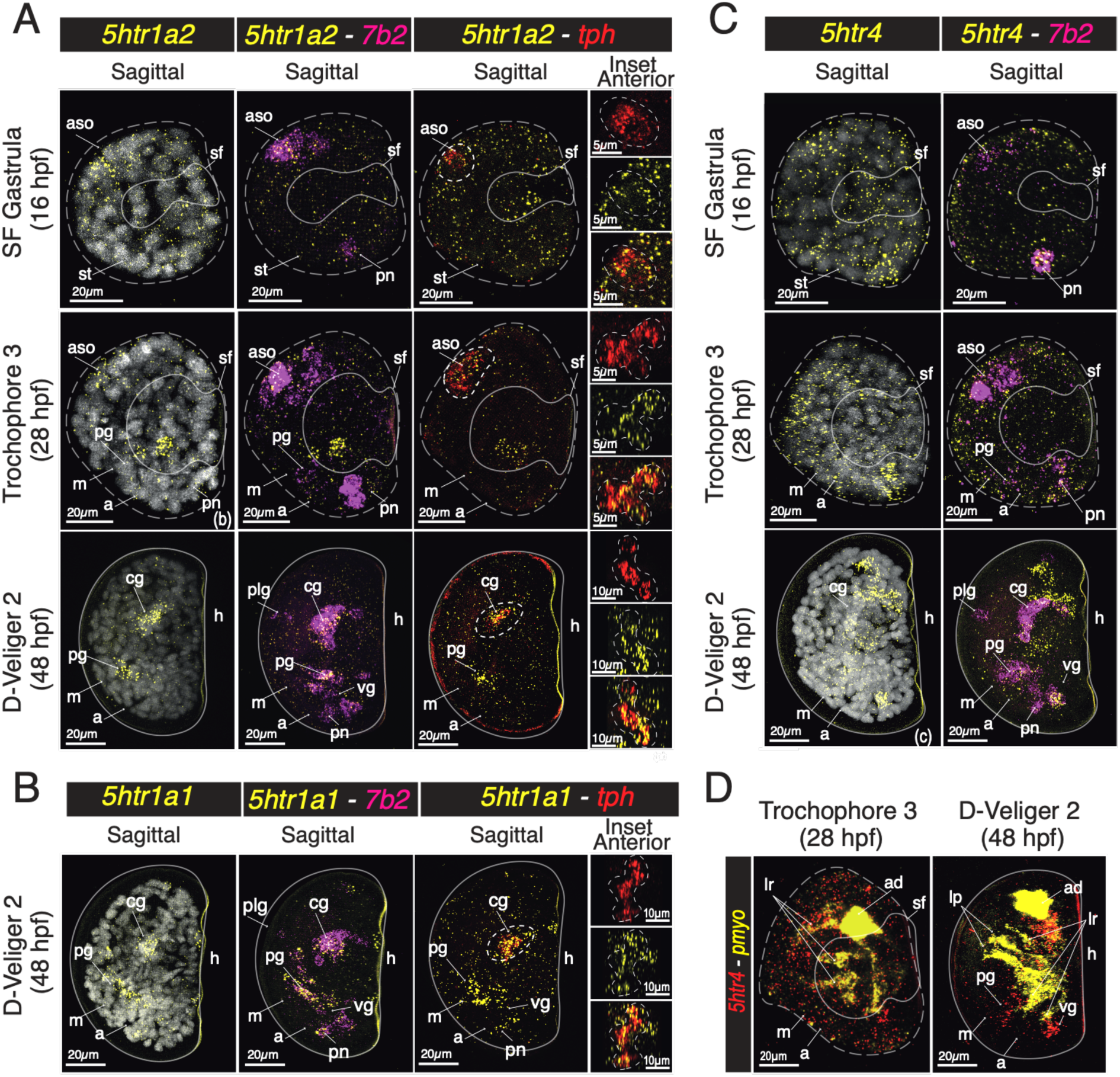
Developmental Expression of serotonin receptors *5-htr 1a1*, *1a2* and *4*. A) Developmental expression of *5-htr1a2* with *7b2* and *tph*. B) Expression of *5-htr1a1* with *7b2* and *tph* in D-Veliger larvae. C) Developmental expression of *5-htr4* with *7b2*. D) Co-localization of *5-htr4* in the muscle system with paramyosin (*pmyo*). In A and B insets display the anterior view of the apical sensory organ (aso) and cerebral ganglion (cg) with the single and multiplexed genes channels. aso: apical sensory organ; m: mouth; a: anus; sf: shellfield; cg: cerebral ganglion; h: hinge; ve: velum; pg: pedal ganglion; plg: pleural ganglion; pn: posterior neurons; vg: visceral ganglion; ad: adductor anterior muscle; lr: larval retractor muscles; pl: larval protractor muscles. Dotted grey lines outline the larval body, continuous grey lines outline the growing shell, dotted white lines outline the areas of co-localization in the aso/cg displayed in the inset. Scalebars as indicated.

**Figure 7.**
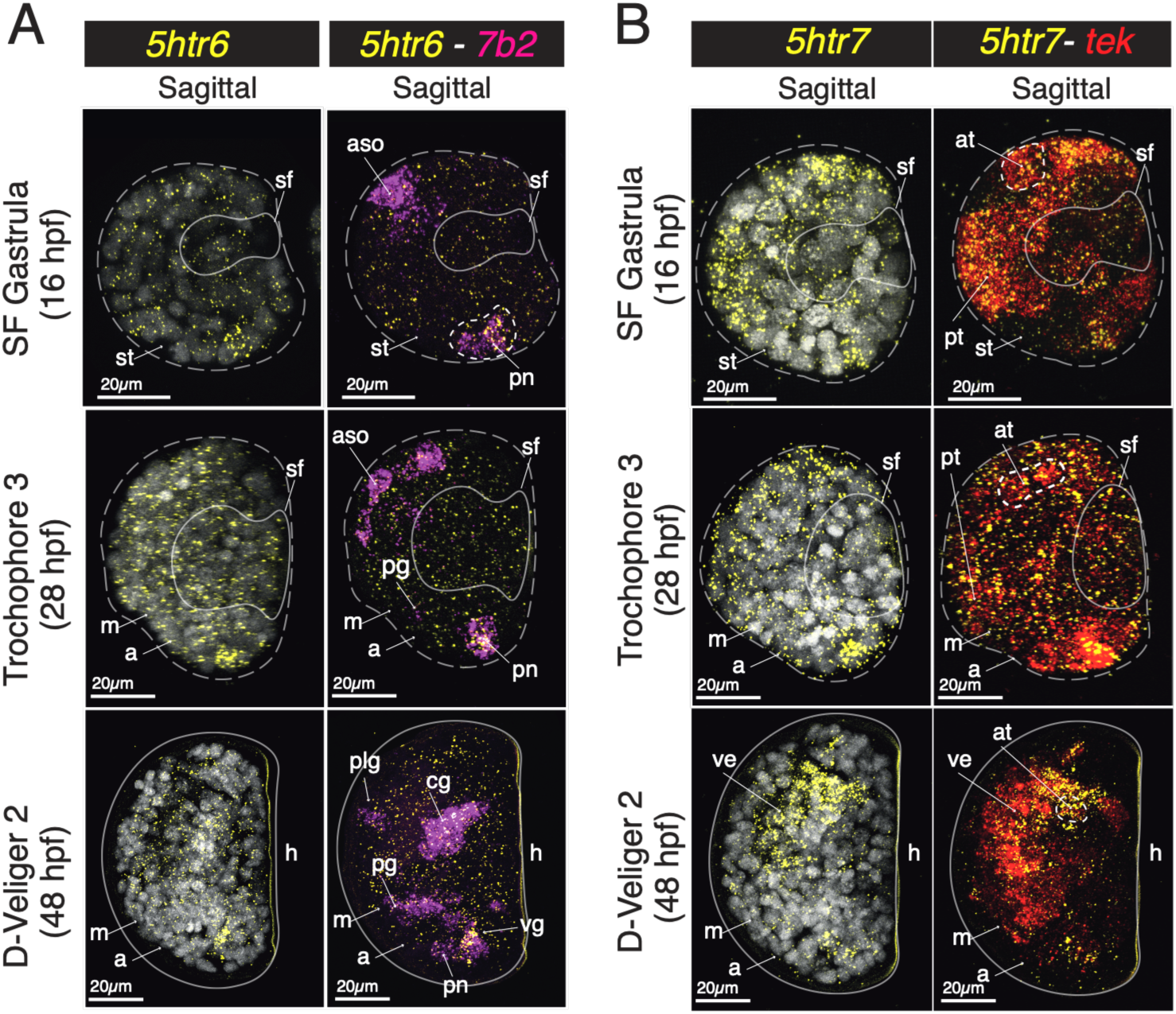
Developmental Expression of 5-htr 6 and 7. A) Developmental expression of *5-htr6* with *7b2*. B) Expression of 5-htr7 with Tektin (*tek*). aso: apical sensory organ; m: mouth; a: anus; sf: shellfield; cg: cerebral ganglion; h: hinge; ve: velum; pg: pedal ganglion; plg: pleural ganglion; pn: posterior neurons; vg: visceral ganglion. Dotted grey lines outline the larval body, continuous grey lines outline the growing shell, dotted white lines outline the areas of the apical tuft. Scalebars as indicated.

### Expression of dopamine receptors

Next, we studied the developmental expression of DA receptors (Fig 8, 9), which were all detectable starting from the trochophore stage. *dr1* displayed expression in the *th*+ cells of the pg (Fig. 8A), in the ciliated epithelium of the prototroch and velum, especially in the area surrounding the mouth (Fig. 8B). *dr2* displayed weak expression in *th*+ cells in both trochophores and D-Veligers, localized in the outer ciliated epithelium (Fig. 8C) as well as in the developing anterior adductor muscle, larval retractors and in two peripheral clusters at the margins of the hinge of D-Veliger larvae (Fig. 8D). On the other hand, *dr3-like* only displayed weak expression in *th*+ cells of the *p*g in D-Veliger larvae and a broader expression in the growing margin of the larval shell and *tph*+ cells (Fig. 8E).

**Figure 8.**
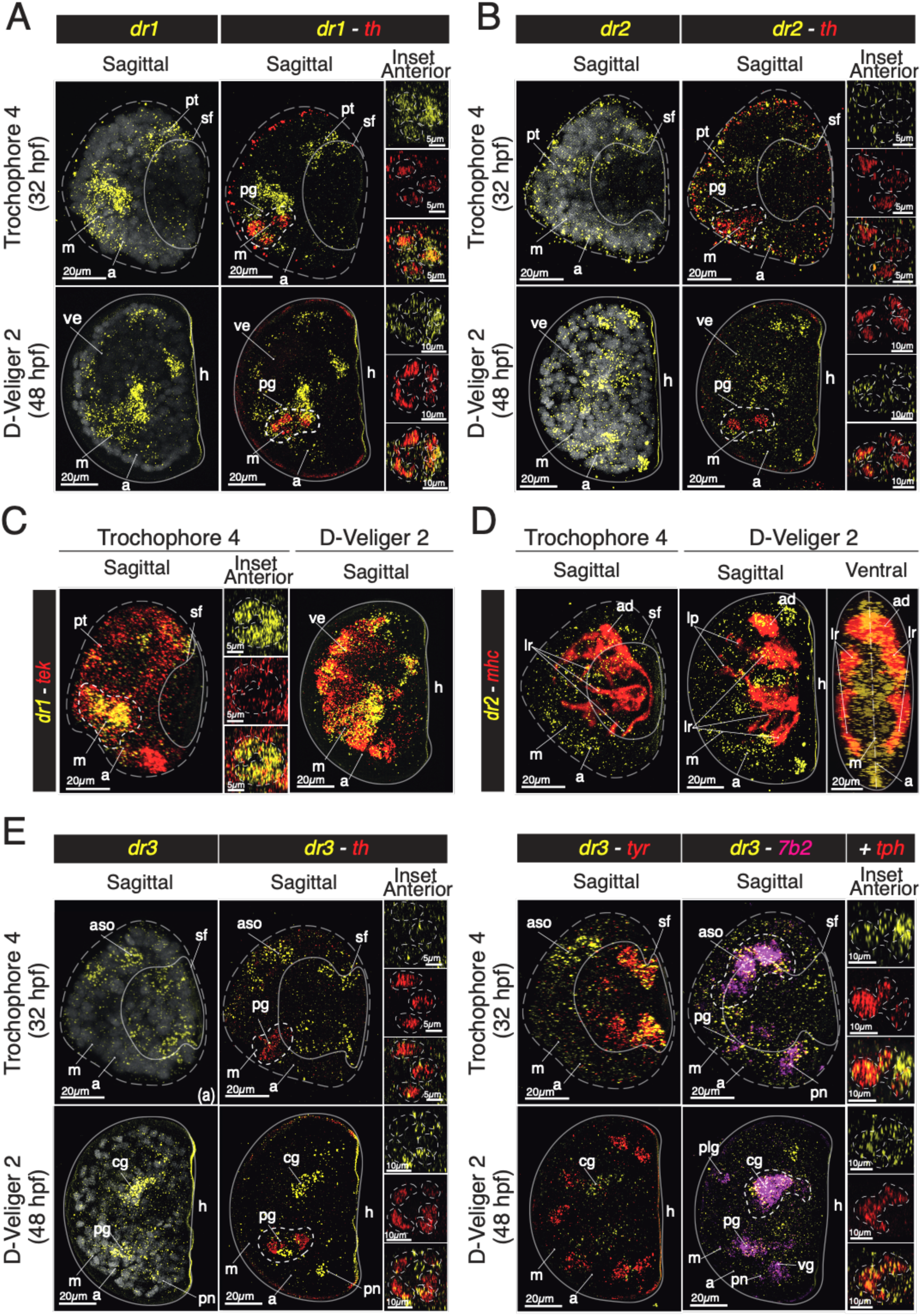
Developmental Expression of dopamine receptors. A) Developmental expression of *dr1* with *th*. B) Expression of *dr1* with *tek*. C) Developmental expression of *dr2* with *th*. D) Developmental expression of *dr2* with *mhc.* E) Developmental expression of *dr3-like* in co-localization with *th*, *tyr*, *7b2* and *tph*. In A and C insets display the anterior view of the pedal ganglion (pg) with the single and multiplexed genes channels. In B insets display the anterior view of the mouth with the single and multiplexed genes channels. In E, insets display the anterior view of the pg with *th* and *dr3-like* and the aso/cg with *dr3-like* and *tph* in single and multiplexed genes channels. aso: apical sensory organ; m: mouth; a: anus; sf: shellfield; cg: cerebral ganglion; h: hinge; ve: velum; pg: pedal ganglion; plg: pleural ganglion; pn: posterior neurons; vg: visceral ganglion; ad: adductor anterior muscle; lr: larval retractor muscles; lp: larval protractor muscles. Dotted grey lines outline the larval body, continuous grey lines outline the growing shell, dotted white lines outline the areas of co-localization in the pg or mouth displayed in the inset. Scalebars as indicated.

**Figure 9.**
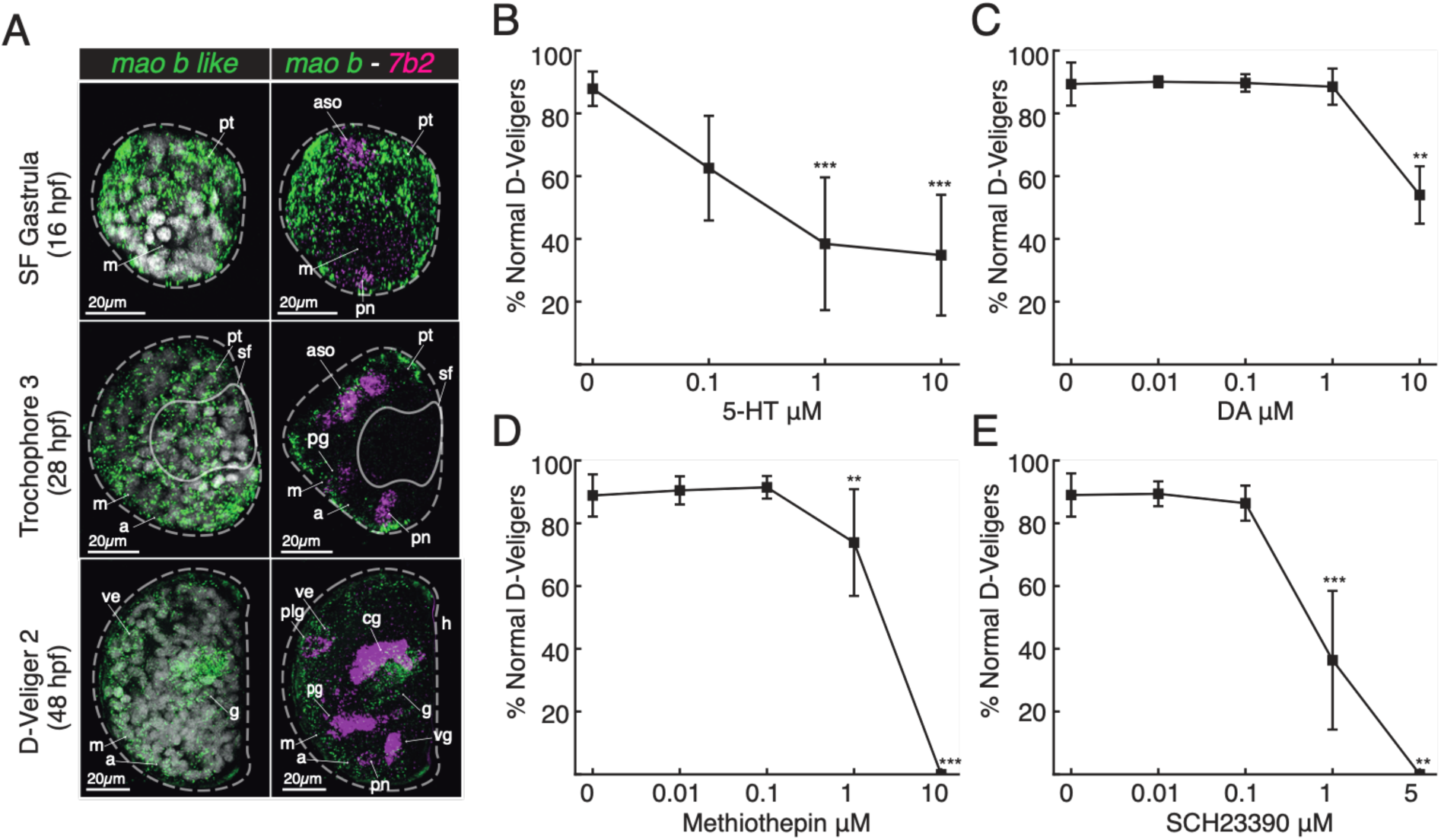
Pharmacological treatments and expression of monoamine oxidase b-like (*mao-b*-like). A-D) Effects of, respectively, serotonin (5-HT), dopamine (DA) and the 5HTRs antagonist Methiothepin and DR1 antagonist SCH23390 on mussel early larval development by standard embryotoxicity assay. **p<0.01, ***p<0.001. E) Developmental expression of *mao-b-like* in mussel larvae alone (green) and in co-localization with *7b2* (magenta). aso: apical sensory organ; m: mouth; a: anus; sf: shellfield; cg: cerebral ganglion; h: hinge; ve: velum; pg: pedal ganglion; plg: pleural ganglion; pn: posterior neurons; vg: visceral ganglion. Dotted grey lines outline the larval body, continuous grey lines outline the growing shell. Scalebars as indicated.

### Expression of monoamine oxidase B

Next, we sought to analyse the expression of monoamine oxidases (MAOs), the principal enzymes involved in monoamine catabolism (Fig. 1B, C; Fig. 9A). We assessed the expression of both mao-a-like and mao-b, but only the mao-b gene showed detectable expression (Fig. 9A). The expression was detected throughout mussel development, with a widespread localization in peripheral tissues in both gastrulae and trochophores. The signal was consistently observed in the ectoderm and ciliated epithelial structures, as well as in metabolically active tissues, such as the developing gut in D-Veliger larvae. In contrast, no clear expression was observed in neurosecretory populations.

### Effect of serotonergic and dopaminergic modulation on larval development

Following the identification and spatial characterization of serotonergic and dopaminergic components during early development, we performed pharmacological assays from fertilized eggs to D-Veliger larvae following standard embryo-toxicity protocols with exogenous monoamines (serotonin and dopamine) and inhibitors of their corresponding receptors (methiothepin and SCH 23390, respectively) (Fig. 9) (34, 35). Serotonin induced a clear dose-dependent impairment of larval development, with a significant reduction in normal D-veliger larvae from 1 μM (−65%, p<0.001), mainly associated with protruding mantle and malformed hinge phenotypes (Fig. 9B). In contrast, dopamine affected larval development only at the highest concentration tested (10 μM) (−42 %, p<0.01) (Fig. 9C). Pharmacological inhibition highlighted the importance of these pathways: methiothepin produced mild effects at 1 μM, but it led to 100% malformed larvae at 10 μM (Fig. 9D); a stronger impact was observed for the D1 receptor antagonist SCH 23390, that significantly reduced normal development from 1 μM (−42%, p<0.01) and caused complete developmental arrest at 5 μM (Fig. 9E) as previously described (34).

## Discussion

In this work, we integrated phylogenetic, transcriptomic, histochemical and pharmacological approaches to provide the first comprehensive characterization of the onset and early organization of the larval monoaminergic system in bivalve molluscs, using the environmentally relevant model system *Mytilus galloprovincialis*.

### A conserved but differentially specialized monoaminergic architecture defines bivalve molluscs

Our comparative analyses revealed that bivalves possess a highly conserved and coherent monoaminergic repertoire, with all analysed genomes displaying a similar complement of monoaminergic components (Fig. 1C). This conservation reflects the retention of an ancestral bilaterian monoaminergic toolkit, although lacking several lineage-specific innovations of vertebrates and ecdysozoans (1, 4, 16, 21, 22, 27). While previous studies in *M. galloprovincialis* and other bivalves reported only a limited subset of monoaminergic elements (23, 24, 57, 66, 70, 71), our results show that up to six monoaminergic circuits can be delineated in pteriomorph bivalves with different degrees of completeness and potential functionality: serotoninergic, dopaminergic, noradrenergic, tyraminergic, octopaminergic and histaminergic. Among these pathways, serotoninergic and dopaminergic systems display a fully conserved and canonical organisation (1, 21, 22). Both retain the complete molecular machinery required for signalling, including synthesis enzymes (*tph/th* and *aadc*), vesicular and membrane transporters (*slc18* and *slc6* families) and multiple receptor subtypes. The retrieved molecular elements are consistent with current models of monoaminergic evolution in Bilateria, including the absence of vertebrate-specific elements such as 5-HTR3, and the retention of invertebrate-specific receptor subtypes such as DR2/inv (1, 21, 22, 66).

In contrast, noradrenergic, tyraminergic and histaminergic systems appear molecularly partial although likely functional. The noradrenergic pathway includes DβH and α-adrenergic receptors but lacks canonical NET transporters and β-adrenergic receptors, suggesting that monoamine clearance may rely on broader-specificity transporters such as OAT (4, 6, 8, 27, 70, 72–75). Similarly, tyraminergic and histaminergic pathways possess synthesis enzymes and receptors but lack dedicated transport systems, consistent with the unresolved nature of monoamine transport in many invertebrate systems (1). These circuits therefore appear functionally integrated within a broader monoaminergic network rather than operating as fully specialised, independent pathways. The octopaminergic system displays a different condition. Although octopamine receptors and transporters are present and the neurotransmitter is detected in the tissues of molluscs, the canonical synthesis enzyme TβH is absent, consistent with its restriction to Ecdysozoa (16, 27, 59, 72, 73, 76–78). Octopamine synthesis in bivalves might therefore rely on alternative enzymatic routes (i.e., DβH) (12, 76) or may occur at reduced levels in favour of tyraminergic signalling. Together, these patterns indicate that monoaminergic circuits in bivalves display different degrees of molecular specialisation depending on whether they rely exclusively on ancestral bilaterian components or on lineage-specific innovations that evolved secondarily in other clades. Overall, the coexistence of complete, partial and incomplete circuits supports the view that monoaminergic signalling in bivalves reflects an early bilaterian condition characterized by molecular versatility and co-option of broad-specificity components.

### Transcriptomic dynamics reveal staged deployment of monoaminergic circuits during development

The analysis of monoaminergic gene expression during embryo–larval development (0–48 hpf) reveals a progressive and staged deployment of monoaminergic circuits during development (Fig. 2, 10A). Genes associated with serotoninergic and dopaminergic signalling, corresponding to the most complete monoaminergic pathways, display sustained and progressively increasing expression throughout larval stages. Serotonergic components peak already in late embryogenesis, whereas dopaminergic components become prominent from the mid-trochophore stages onwards. As development proceeds, both systems progressively acquire additional molecular complexity through the expression of multiple receptor subtypes, transporters and catabolic enzymes. In contrast, partial or incomplete circuits display more restricted temporal profiles. Tyraminergic components are predominantly associated with early embryogenesis; noradrenergic signalling emerges mainly during veliger stages, whereas histaminergic and octopaminergic are associated to both embryogenesis and, more strongly, to late larval development. These temporal differences suggest that monoaminergic circuits contribute differently to successive phases of development, with serotoninergic and dopaminergic pathways supporting sustained developmental regulation while other circuits likely perform more restricted or stage-specific functions. Importantly, several receptors and transporters are expressed prior to the appearance of their corresponding synthesis enzymes. Serotonin receptors such as *5-htr1a1/2*, *5-htr6*-like and *5-htr7*, as well as dopaminergic components including *dr2*, *dr3-like-*like and *idat,* are detected before *tph* or *th* expression (Fig. 2). Similarly, maternal expression of *aadc* in unfertilised eggs suggests an early capacity for monoamine production. Together, these observations indicate that early monoaminergic signalling may initially depend on maternal or environmental monoamines before the establishment of endogenous synthesis (1, 2, 23, 24, 79).

This organisation is consistent with the ancestral developmental functions of monoamines described across Bilateria, where serotonin and dopamine act as pre-neuronal regulators of cleavage, morphogenesis, cytoskeletal dynamics and gene expression (1, 2, 23, 24, 79, 80). In this context, monoaminergic signalling may operate as an interface between intrinsic developmental programs and external environmental conditions. The widespread occurrence of indole-derived compounds in marine environments (81–83), together with the early expression of receptor repertoires in embryos and larvae, further supports the possibility that developing larvae integrate environmental and endogenous monoaminergic cues simultaneously to regulate distinct developmental processes (23, 46, 80, 84–87). Altogether, these data support a developmental model in which monoaminergic signalling is progressively deployed during embryogenesis and larval differentiation, with increasing molecular complexity accompanying tissue regionalisation and functional maturation (23, 46, 84–87).

### A distributed neurosecretory scaffold precedes ganglionic organisation in bivalve larvae

To establish the neuroanatomical framework within which monoaminergic signalling operates, we characterised the development of the neurosecretory system using the neuroendocrine marker *7b2* (57, 68). The observed expression patterns indicate that the larval ns system of *M. galloprovincialis* initially develops as a peripheral network rather than as a ganglionic system. At the end of gastrulation, *7b2*+ neurosecretory cells are already regionalised into anterior and posterior populations associated with the presumptive apical sensory organ and posterior apical territories (41–44, 88, 89). This organisation demonstrates that neurosecretory differentiation precedes the establishment of definitive ganglia and is consistent with recent models of molluscan neurogenesis, in which peripheral sensory and neurosecretory populations emerge prior to ganglionic structures (42, 45, 69). During trochophore development, the expansion of anterior neurosecretory territories together with the emergence of ventral populations reflect progressive organization of the nervous system along the anterior–posterior and dorsoventral axes (41–44, 88, 89). By the D-veliger stage, *7b2*+ territories correspond to the presumptive cerebral, pedal, pleural and visceral ganglia, indicating that the definitive ganglionic architecture develops through regionalisation and integration of this neurosecretory scaffold (41–44, 88, 89). The close association of *7b2*+ cells with the apical tuft, ciliated epithelia and larval musculature further suggests that these early neurosecretory populations integrate sensory inputs with physiological and behavioural responses before the emergence of mature neural circuits (42, 44, 45, 69). Together, these findings support the view that the larval nervous system of bivalves initially functions as a distributed sensory-secretory network that progressively acquires ganglionic structure and functional specialisation (89).

### Serotoninergic and dopaminergic systems are spatially segregated but functionally interconnected

Based on the developmental expression analyses, *in situ* HCR focused on the serotonergic and dopaminergic systems, which represent the only monoaminergic pathways with complete molecular repertoires and sustained expression throughout embryo–larval development. The localization of biosynthetic enzymes, transporters and vesicular machinery resolved these pathways into two distinct neurosecretory circuits. The serotonergic system emerges first, with *tph*, *aadc*, *vmat* and later *sert1/2* defining a functional serotonergic domain centred on the aso/cg from late gastrulation onwards. While transmitter synthesis and vesicular loading remain restricted to this anterior territory, serotonin transporters display broader distributions, particularly *sert2* in ciliated epithelia in addition to lateral serotonergic neurons (Fig. 10B). This spatial segregation suggests that serotonin signalling extends beyond sites of synthesis and may operate through both local synaptic and broader paracrine mechanisms (3, 8, 23, 24, 36, 46, 79, 84–87, 90–94). In contrast, the dopaminergic system appears later and is associated with ventral neurosecretory territories corresponding to the developing pedal ganglion (9, 10, 25, 44, 80, 95). The coordinated expression of *th, aadc*, *vmat* and *idat* supports the establishment of an active dopaminergic circuit linked to the ventral axis and oral region (Fig. 10C). Notably, *idat* expression in muscles and other non-neuronal tissues indicates that dopaminergic signalling is not restricted to neuronal populations, supporting broader neuroeffector functions during larval development (9, 16, 24, 66, 80). Together, these observations reveal two spatially segregated monoaminergic circuits: an early anterior serotonergic one associated with sensory and ciliated territories and a later ventral dopaminergic one associated with oral and muscular domains. Their organization mirrors the major body axes of the larval nervous system, as well as previous knowledge of the physiological actions of the neurotransmitters in adult molluscs and suggests that monoaminergic signalling is incorporated into the neurosecretory scaffold from which the definitive ganglionic system develops (23, 25, 42, 45, 69, 89). The distribution of monoaminergic receptors further refines this organization by revealing signalling domains that extend beyond sites of monoamine synthesis. Within the serotonergic system, some receptors colocalise with *tph*-expressing neurons of the aso/cg, whereas others are associated with posterior, visceral and ciliated territories as well as the muscle system. Likewise, dopaminergic receptors display broader distributions than the *th*+ pedal domain, with expression detected in epithelial, muscular and shell-forming tissues (Fig. 10D). As previously documented in adults (2, 23–25), these patterns indicate that monoaminergic signalling is not restricted to transmitter-producing neurons but operates across multiple larval compartments through both neuronal and non-neuronal mechanisms already in developing larvae (3, 8, 16, 44, 66, 71, 80, 93, 96, 97). Importantly, our data show overlaps between distinct monoaminergic circuits. In particular, the presence of *dr3-like*-like within *tph*-expressing regions and 5-HTR1A1/2 within th+ cells/pg suggests potential interactions between serotonergic and dopaminergic pathways. Comparable reciprocal regulation between these systems is widely documented in vertebrates, for example in raphe nuclei, where dopaminergic signalling modulates serotonergic activity (95, 98). Although the functional significance of these interactions remains unresolved in bivalves, the observed overlap suggests that monoaminergic pathways do not operate as isolated modules but rather form an integrated signalling network within the developing neurosecretory system. Taken together, the spatial distribution of biosynthetic enzymes, transporters and receptors indicates that monoaminergic circuits emerge within the same neurosecretory territories that give rise to the definitive ganglionic system. Rather than representing a modulatory layer added onto an already established nervous system, monoaminergic signalling appears integrated into the neurosecretory scaffold early in neural development. The progressive appearance of monoaminergic components across neuronal and non-neuronal tissues further suggests that these circuits are assembled alongside the developing nervous system itself, positioning them as potential mediators between developmental programs and environmental inputs (1, 2).

**Figure 10.**
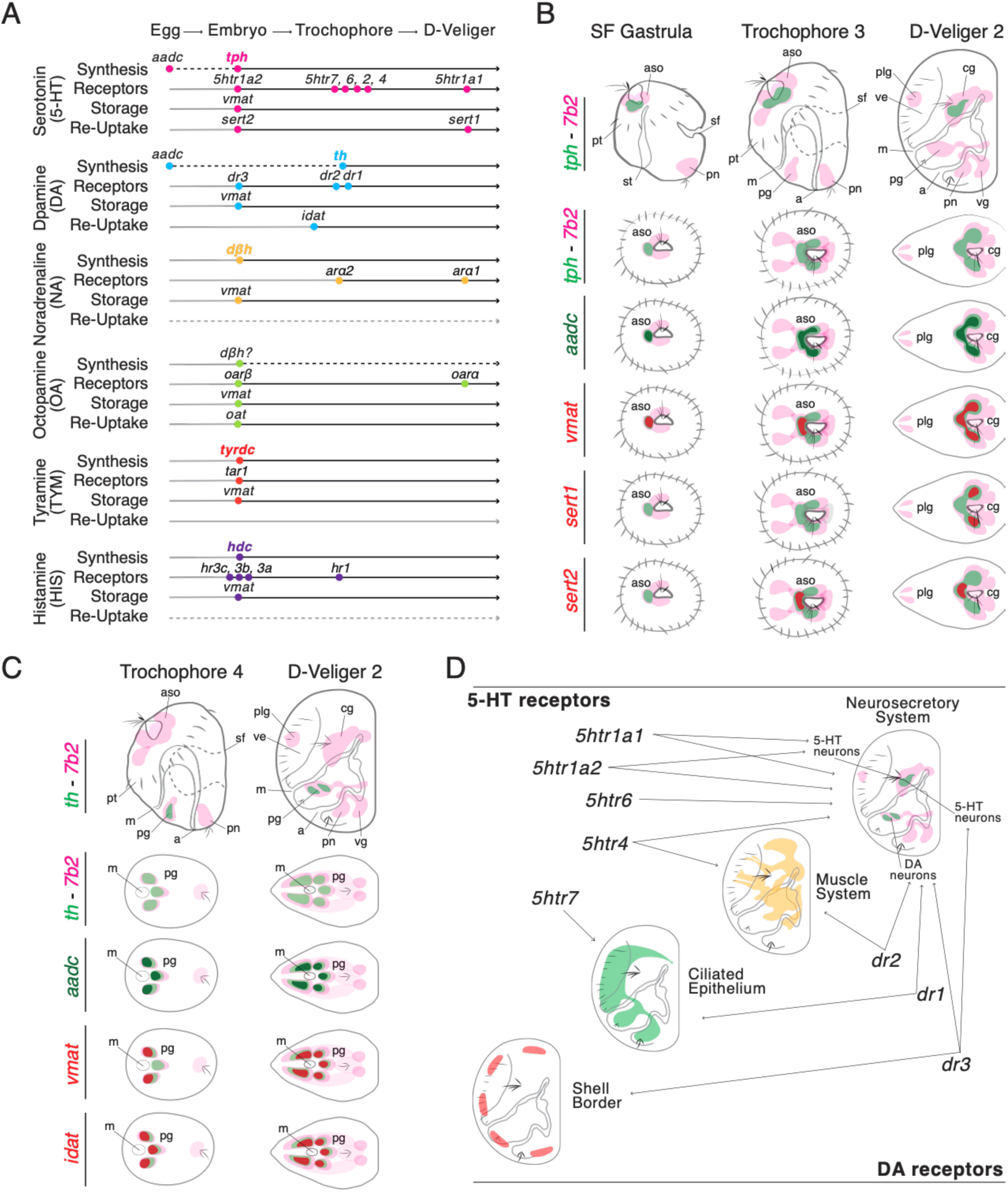
The onset of Monoaminergic system signalling in the Mediterranean mussel. A) Schematic representation of the temporal deployment of the six monoaminergic circuit identified in *M. galloprovincialis* larvae. For each circuit, dots indicate first detectable expression (TPM > 0), black lines indicate time course of expression, dotted lines indicate undetected specific elements, rate limiting enzymes are highlighted in the corresponding circuit colours. B) Schematic representation of the onset of serotoninergic pre-synaptic machinery. C) Schematic representation of the onset of the dopaminergic pre-synaptic machinery. D) Schematic representation of the expression patterns of the serotoninergic (*5htr*) and dopaminergic (*dr*) receptors analysed by *in situ* HCR in the study.

### Pharmacological validation of monoaminergic circuits hints at distinct roles of serotonin and dopamine in bivalve development

Pharmacological perturbation confirmed the functional relevance of serotonergic and dopaminergic signalling during early development (Fig. 9). Consistent with the developmental expression profiles and spatial distribution of monoaminergic components, 5-HT exposure produced strong dose-dependent developmental defects, whereas DA elicited comparatively weaker responses (23, 24, 35, 71, 80). In addition, inhibition of 5HTRs with methiothepin induced arrested trochophores suggesting an early effect in development that could be due to a non-neuronal role of 5HT in *Mytilus* early development. Further research is required to assess whether this role of serotonin signalling is comparable to the ancestral morphogenetic role of serotonin found in *Drosophila*, *Xenopus* and chicken embryos (99, 100).

Despite a low sensitivity to micromolar concentrations of DA, inhibition of dopaminergic receptors using SCH23390 caused severe developmental impairment at relatively low concentrations, supporting an essential role of DA signalling during larval development (23, 24, 35, 71, 80). These findings are broadly consistent with previous studies in bivalves and other invertebrates showing that monoaminergic pathways contribute to larval growth, shell formation and developmental regulation (23, 24, 35, 71, 80). The expression of *dr3-like* around the shell field at 48 hpf suggests that this receptor would be the one regulating shell morphogenesis and might underpin the strong sensitivity of shell morphogenesis to environmental insults (30, 33, 34). The comparatively high concentrations of 5HT/DA required to elicit pharmacological effects may reflect the activity of endogenous buffering mechanisms. In particular, the expression of *mao-b* in epidermal and metabolically active tissues suggests that monoamine catabolism may already be active during larval stages, limiting the accumulation of exogenous compounds (1, 20, 64). Together, these results provide functional support for the developmental relevance of monoaminergic signalling and complement the molecular and anatomical evidence presented here.

## Conclusions

Our results demonstrate that the bivalve monoaminergic system comprises up to six circuits representing an ancestral bilaterian monoaminergic toolkit. This toolkit retains the core components of the signalling network while lacking the lineage-specific innovations that emerged in vertebrates and ecdysozoans. Monoaminergic pathways are established early in embryogenesis and are progressively assembled alongside the developing nervous system, with serotonergic and dopaminergic circuits emerging as the predominant signalling systems. The early expression of receptors and transporters, often preceding endogenous monoamine synthesis, together with their localization in both neuronal and non-neuronal tissues, supports developmental and regulatory functions that extend beyond canonical neurotransmission.

By integrating phylogenetic, transcriptomic and anatomical evidence, this study provides a comprehensive framework for understanding the assembly and diversification of monoaminergic systems in Spiralia. These findings reinforce the view that monoaminergic signalling is an ancient and deeply conserved regulatory network linking development, neuroendocrine function and environmental responsiveness across Bilateria. Future studies should resolve the onset of embryonic histaminergic and tyraminergic circuits to establish a more complete picture of the temporal deployment and developmental contributions of individual monoamines. Combined with receptor deorphanization and analytical approaches that quantify monoamine synthesis, transport and clearance through development in a comparative framework, such work will define the kinetics of monoaminergic signalling and clarify both its conserved and lineage-specific functions in developmental physiology, environmental plasticity, and the evolution of nervous systems and species resilience to global change and pollution.

## Ethics approval and consent to participate

Ethics approval was not required for this work. Consent to participate was not required for this work.

## Consent for publication

Not applicable.

## Availability of data and materials

The RNAseq data analysed during the current study are available in the NCBI SRA database under accession number PRJNA996031. Image datasets, including negative results, will be made available upon request to the authors.

## Funding

This study was funded by: Centre National de la Recherche Scientifique DBM 2021 program and the Agence Nationale de la Recherche (ANR-21-CE34–0006–02 - R. Dumollard); Fondi Ricerca Ateneo2023, Università degli Studi di Genova (T. Balbi); PhD in Marine Sciences (Marine Biology and Ecology), 37th Cycle (100,022–2022-ALTRIPOSTL-RISSO), Università degli Studi di Genova; Consolidator fellowship 2022 (Università degli studi di Genova, A. Miglioli).

### Author’s Contributions

**BR:** Writing – review & editing, Visualization, Methodology, Investigation, Funding acquisition, Formal analysis, Data curation. **JB, LB, TB:** Investigation, Data curation, review and editing. **RD:** Writing – review & editing, Writing – original draft, Visualization, Supervision, Resources, Methodology, Investigation, Funding acquisition, Data curation, Conceptualization. **LC:** Writing – review & editing, Writing – original draft, Visualization, Supervision, Methodology, Investigation, Data curation, Conceptualization, Funding acquisition. **AM:** Writing – review & editing, Writing – original draft, Visualization, Methodology, Investigation, Formal analysis, Data curation, Conceptualization, Funding acquisition.

## Acknowledgements

The authors would like to express their appreciation for the Laboratoire de Biologie du Développement de Villefranche-sur-Mer (LBDV), Villefranche-sur-Mer, and to thank Sébastien Schaub of the Plateforme d’Imagerie par Microscopie (PIM) as well as Laurent Gilletta and Axel Duchene of the Service Aquariologie (SA) of the Institut de la Mer de Villefranche (IMEV), France, which is supported by EMBRC-France. The research leading to these results has been conceived under the PhD Course in Marine Science and Technologies (MST), Curriculum: Marine Ecosystem Science, XXXVIII cycle, University of Genoa, Italy (https://corsi.unige.it/en/corsi/11539). This study represents partial fulfilment of the requirements for the PhD thesis of Beatrice Risso at the MST course in co-tutoring with the Sorbonne University. The authors would also like to thank Dr Kévin Drouet, Dr José-Maria Martín-Duran and Lily Nova Winkler, for the fruitful discussions.

## Supplementary Figures and Legends

**Supplementary Figure 1.**
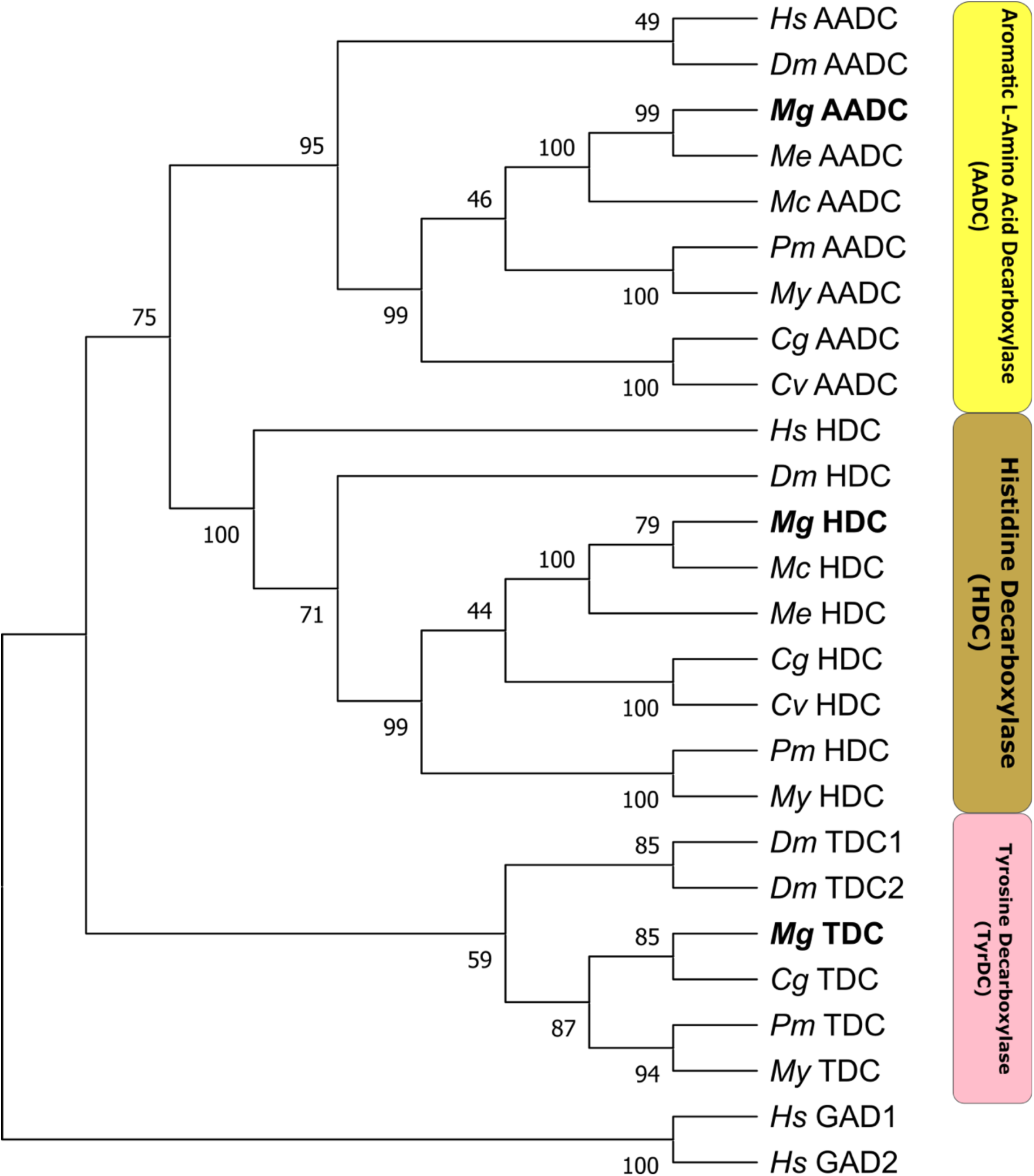
Orthology sequence analysis of decarboxylase proteins: AADC, HDC, and TDC in *M. galloprovincialis* using human and fruit fly proteins as references sequences. The human Glutamate Decarboxylases 1 and 2 (GAD1, GAD2) were used as outgroup. Clades are coloured as follows: AADC in yellow, HDC in brown, and TyrDC (abb. TDC) in rose. *M. galloprovincialis* sequences are highlighted in bold. Sequences from bivalve species with publicly available genome repositories in NCBI were included (*M. edulis, M. coruscus, Magallana gigas, Crassostra virginica, Mizohopecten. yessoensis* and *Pecten maximus*) as previously shown in Canesi et al. (2022). Abbreviations of species and enzymes are listed in Supplementary Table 3. The tree was realised by using Maximum Likehood method and Le Gascuel model; bootstraps values were inferred from 1000 replicates. Initial tree(s) for the heuristic search were obtained automatically by applying Neighbor-Join and BioNJ algorithms to a matrix of pairwise distances estimated using the JTT model, and then selecting the topology with superior log likelihood value. A discrete Gamma distribution was used to model evolutionary rate differences among sites (5 categories (+G, parameter = 1.0119)). This analysis involved 26 amino acid sequences. All positions with less than 85% site coverage were eliminated.

**Supplementary Figure 2.**
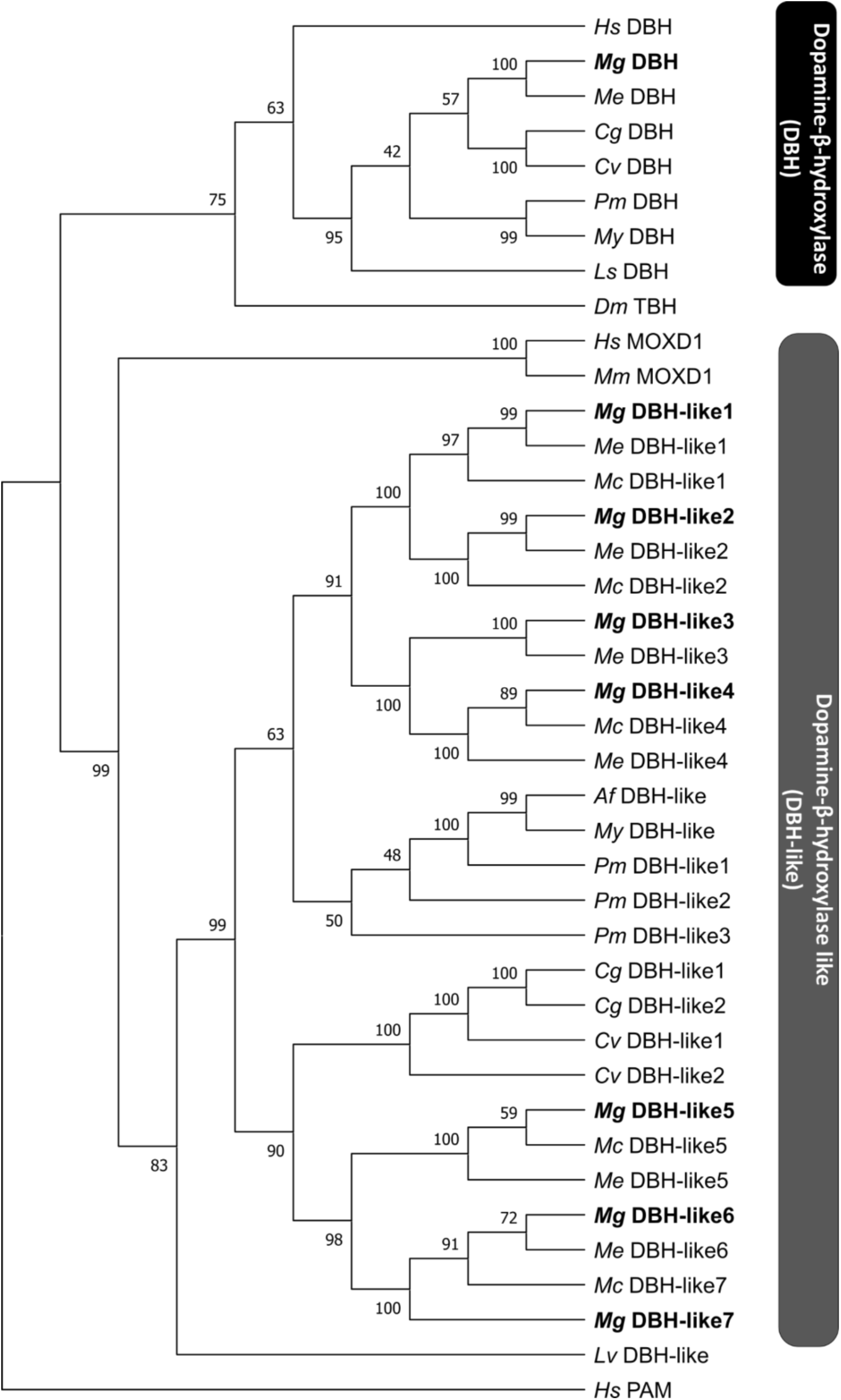
Orthology sequence analysis of of copper-containing hydroxylase proteins: DβH, and DβH-like in *M. galloprovincialis* using human, fruit fly, great pond snail (*Lymnaea stagnalis*), whiteleg shrimp (*Litopenaeus vannamei*), and *Azumapecten farreri* as reference sequences The human Peptidylglycine *α*-amidating monooxygenase (PAM) was used as outgroup. Clades are coloured as follows: DβH in black, and DβH-like in grey. *M. galloprovincialis* sequences are highlighted in bold. Other bivalve species were included as previously described in the methods section. Abbreviations of species and enzymes are listed in Supplementary Table 4. The tree was realised by using Maximum Likehood method and Le Gascuel model; bootstraps values were inferred from 1000 replicates. Initial trees for the heuristic search were obtained automatically by applying Neighbor-Join and BioNJ algorithms to a matrix of pairwise distances estimated using the JTT model, and then selecting the topology with superior log likelihood value. A discrete Gamma distribution was used to model evolutionary rate differences among sites (5 categories (+G, parameter = 1.2529)). The rate variation model allowed for some sites to be evolutionarily invariable ([+I], 0.56% sites). This analysis involved 40 amino acid sequences. All positions with less than 85% site coverage were eliminate. Dopamine-β-Hydroxylase (DBH) is highlighted in black, Dopamine-β-Hydroxylase-like (DBH-like) in dark grey.

**Supplementary Figure 3.**
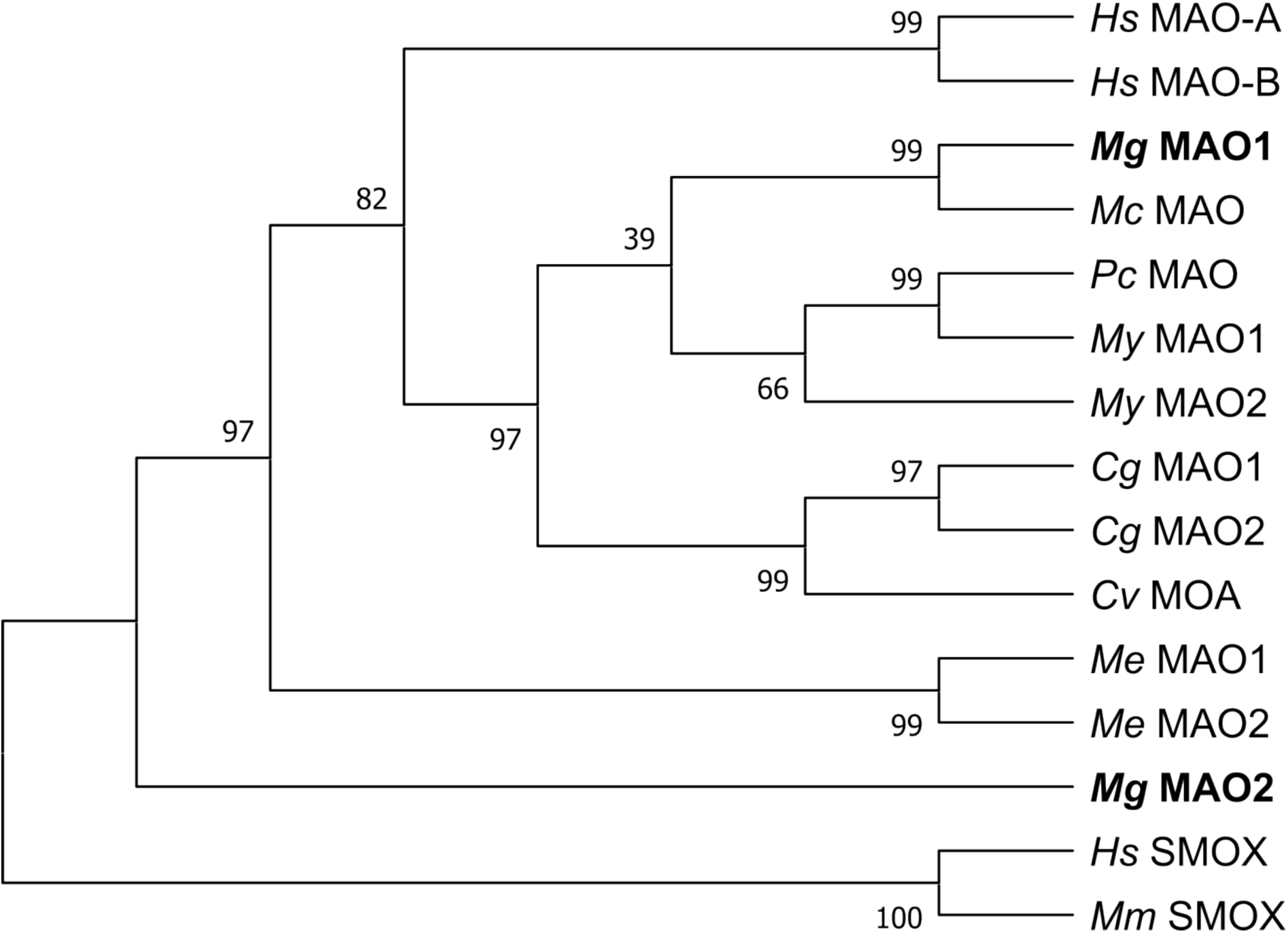
Orthology sequence analysis of MAO proteins in *M.* gallopronvicialis, using human sequences as reference. Human and murine spermidine oxidases (SMOXs) were used as outgroup. *M. galloprovincialis* sequences are highlighted in bold.Abbreviations of species and enzymes are listed in Supplementary Table 5. The tree was realised by using Maximum Likehood method and Le Gascuel model; bootstraps values were inferred from 1000 replicates. Initial tree(s) for the heuristic search were obtained automatically by applying Neighbor-Join and BioNJ algorithms to a matrix of pairwise distances estimated using the JTT model and then selecting the topology with superior log likelihood value. A discrete Gamma distribution was used to model evolutionary rate differences among sites (5 categories (+G, parameter = 1.1999)). This analysis involved 15 amino acid sequences. All positions with less than 85% site coverage were eliminated.

**Supplementary Figure 4.**
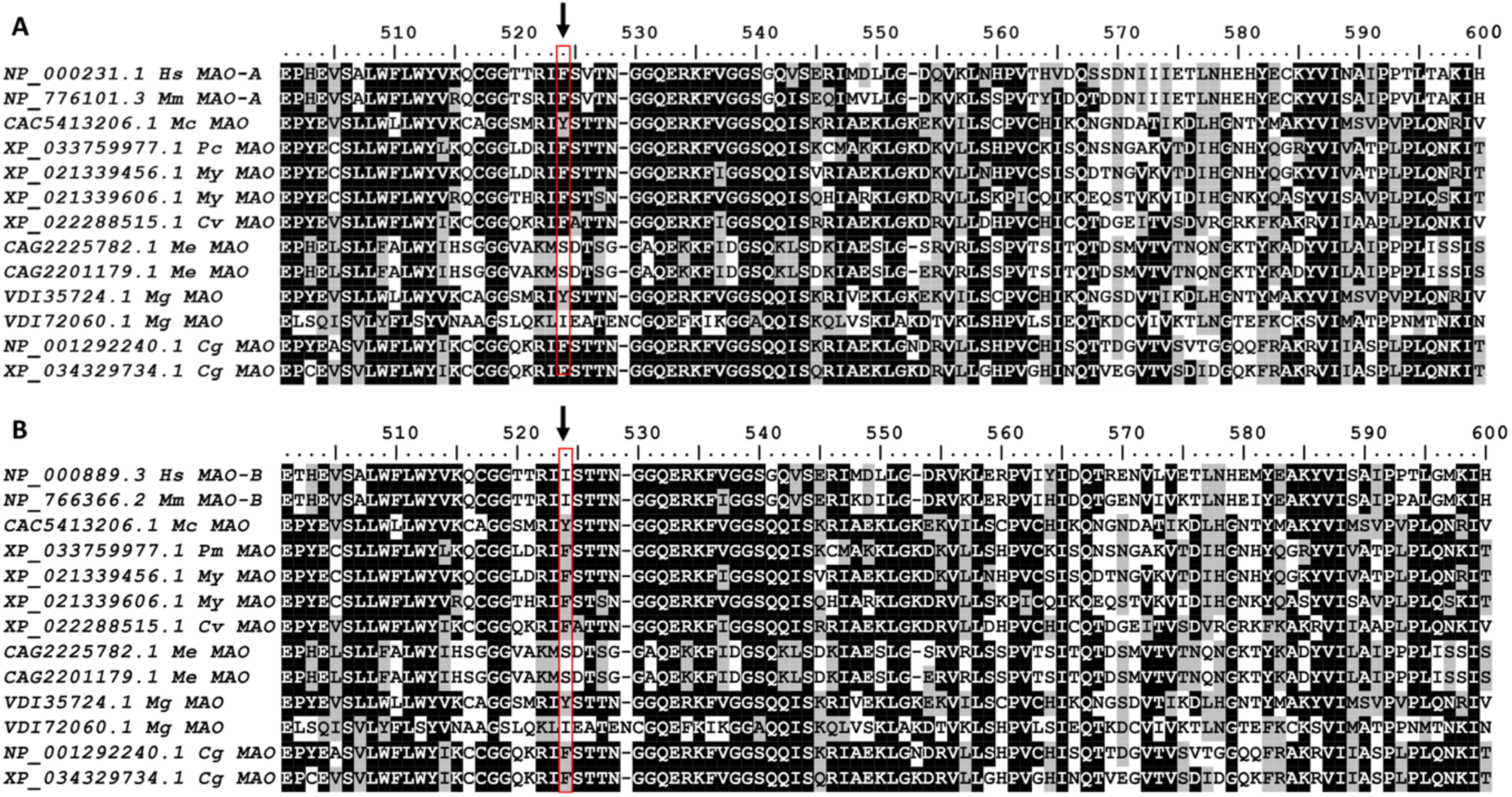
Insight of amino acid multiple sequences alignment of MAOs identified in *M. galloprovincialis* and other bivalve species as described in the methods section. A-B. Sequences alignment with human and murine MAO-A and MAO-B respectively. Black arrow and red box indicate F208 and I199. Black shading indicates identical amino acids, grey shading similar amino acids, absence of shading indicates that amino acids are not conserved. Abbreviations of species and enzymes can be found in Supplementary Table 6.

**Supplementary Figure 5.**
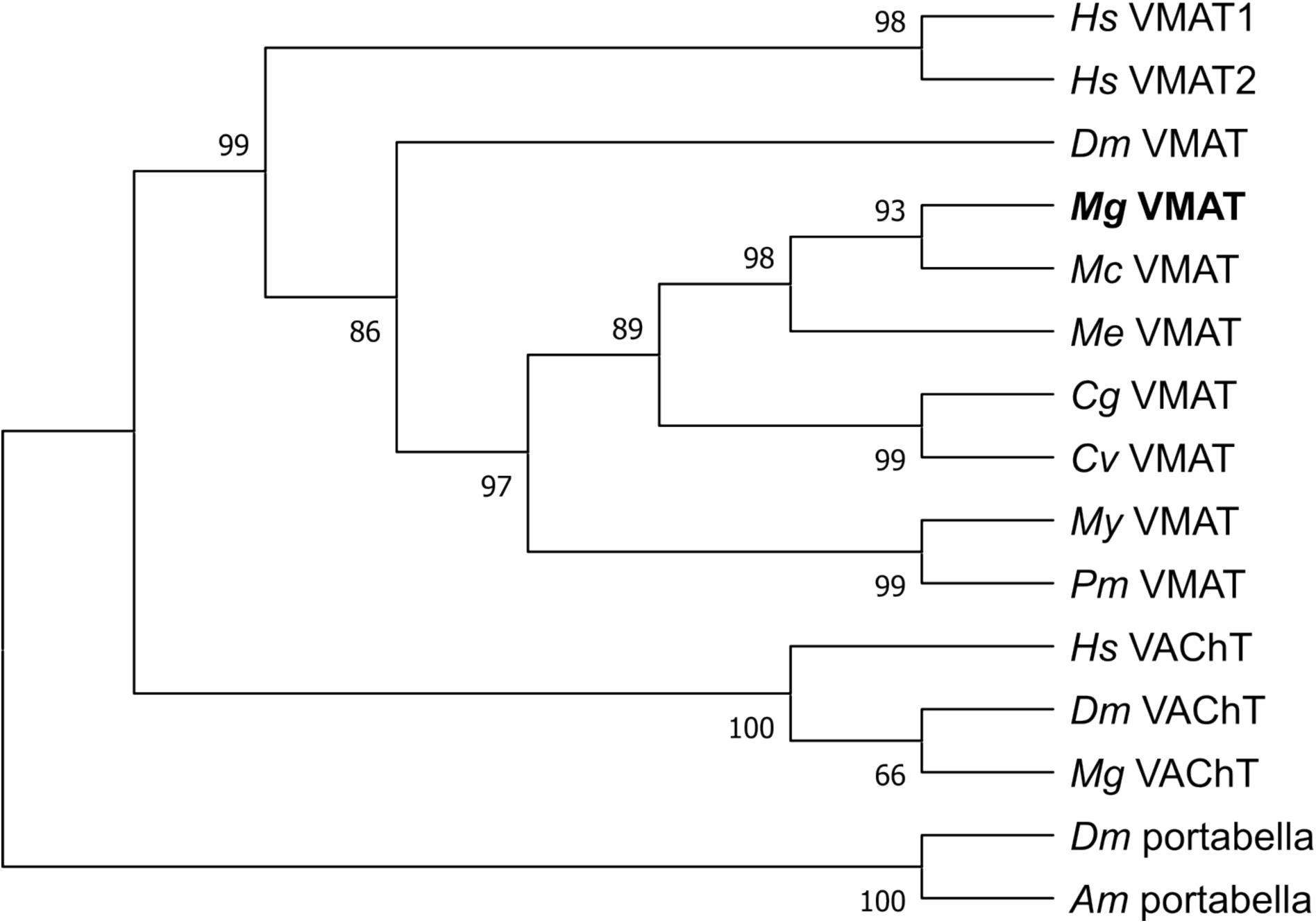
Orthology sequence analysis of VMAT proteins in *M. galloprovincialis* using human and fruit fly sequences as references. Fruit fly and Western honey bee (*Apis mellifera*) orphan vesicular neurotransmitter transporter (portabella) were used as outgroup. *M. galloprovincialis* sequence is highlighted in bold. Other bivalve species were included as previously described in the methods section. Abbreviations of species and enzymes are listed in Supplementary Table 7. The tree was realised by using Maximum Likehood method and Le Gascuel model; bootstraps values were inferred from 1000 replicates. Initial tree(s) for the heuristic search were obtained automatically by applying Neighbor-Join and BioNJ algorithms to a matrix of pairwise distances estimated using the JTT model, and then selecting the topology with superior log likelihood value. A discrete Gamma distribution was used to model evolutionary rate differences among sites (5 categories (+G, parameter = 0.9514)). This analysis involved 15 amino acid sequences. All positions with less than 85% site coverage were eliminated.

**Supplementary Figure 6.**
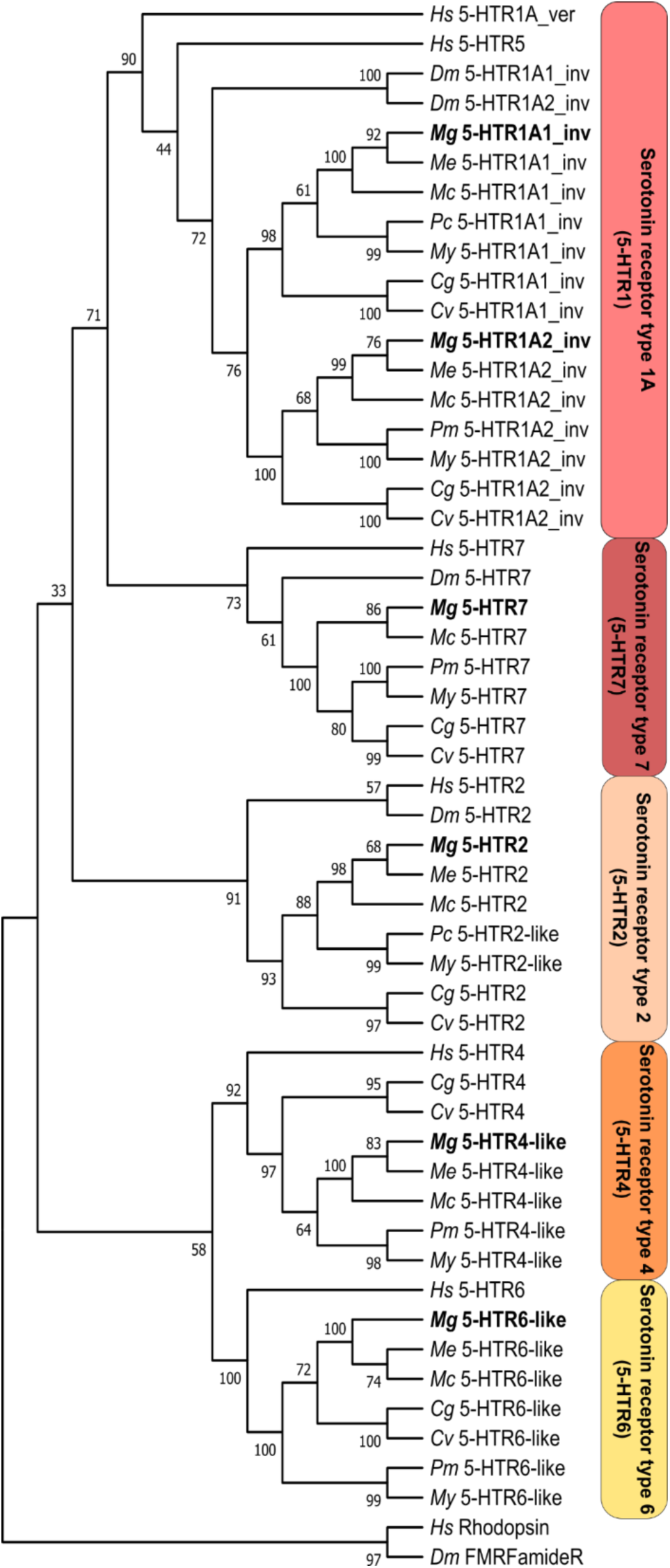
Orthology sequence analysis of 5-HTRs proteins in *M. galloprovincialis* using human and fruit fly sequences as references. Human rhodopsin, and fruit fly FMRFamide receptors were used as outgroup. 5-HTRs subtypes are coloured with different shades of red. *M. galloprovincialis* sequence is highlighted in bold. Other bivalve species were included as previously described in the methods section. Abbreviations of species and enzymes are listed in Supplementary Table 8. The tree was realised by using Maximum Likehood method and Le Gascuel model; bootstraps values were inferred from 1000 replicates. Initial tree(s) for the heuristic search were obtained automatically by applying Neighbor-Join and BioNJ algorithms to a matrix of pairwise distances estimated using the JTT model and then selecting the topology with superior log likelihood value. A discrete Gamma distribution was used to model evolutionary rate differences among sites (5 categories (+G, parameter = 0.8278)). The rate variation model allowed for some sites to be evolutionarily invariable ([+I], 0.66% sites). This analysis involved 59 amino acid sequences. All positions with less than 85% site coverage were eliminated.

**Supplementary Figure 7.**
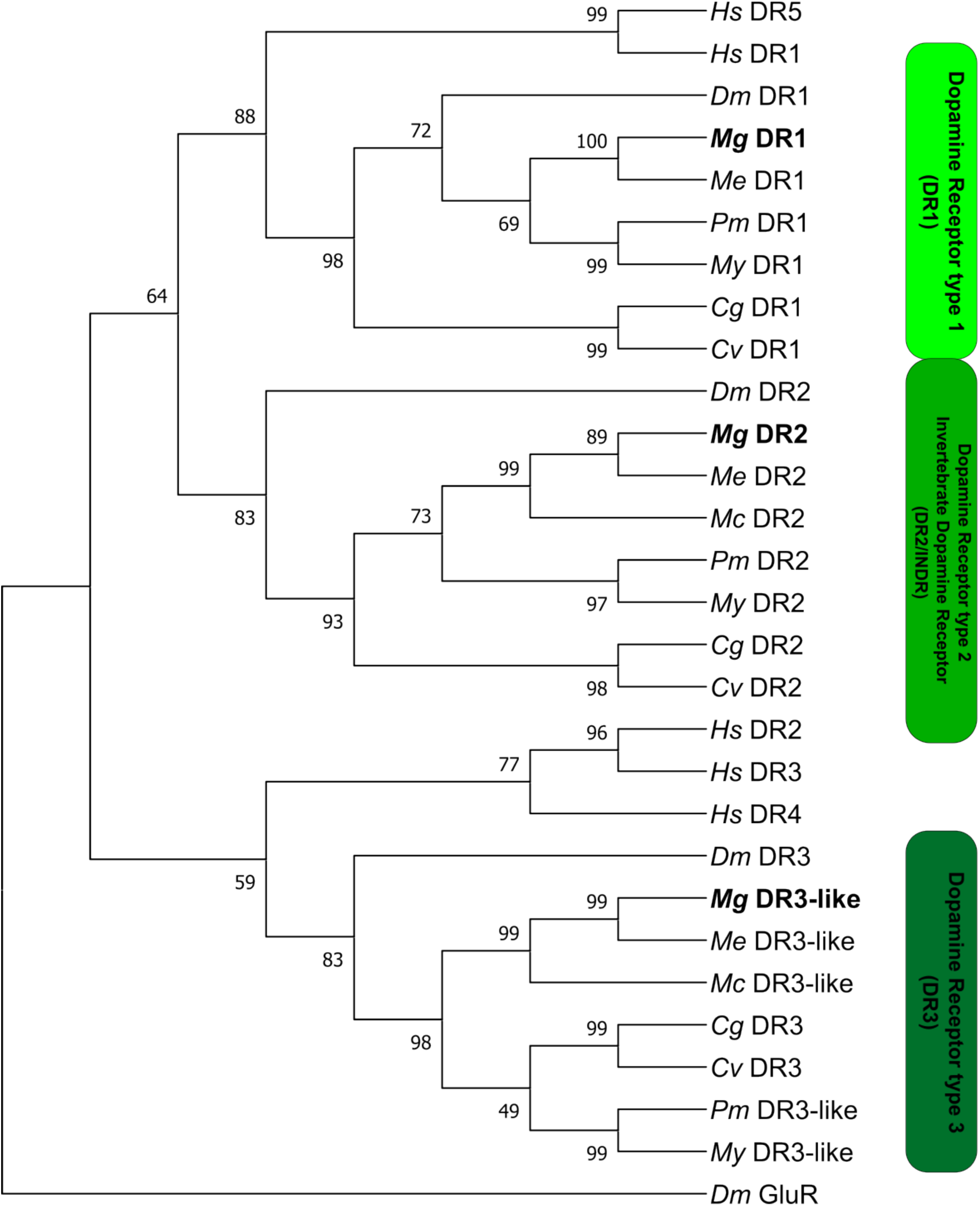
Orthology sequence analysis of DRs proteins in *M. galloprovincialis* using human and fruit fly sequences as references. Fruit fly metabotropic glutamate receptor (GluR) was used as outgroup. DRs subtypes are coloured with different shades of green. *M. galloprovincialis* sequence is highlighted in bold. Other bivalve species were included as previously described in the methods section. Abbreviations of species and enzymes are listed in Supplementary Table 9. The tree was realised by using Maximum Likehood method and Le Gascuel model; bootstraps values were inferred from 1000 replicates. Initial tree(s) for the heuristic search were obtained automatically by applying Neighbor-Join and BioNJ algorithms to a matrix of pairwise distances estimated using the JTT model, and then selecting the topology with superior log likelihood value. A discrete Gamma distribution was used to model evolutionary rate differences among sites (5 categories (+G, parameter = 2.3432)). This analysis involved 29 amino acid sequences. All positions with less than 85% site coverage were eliminated. DRs types are highlighted by different shades of green.

**Supplementary Figure 8.**
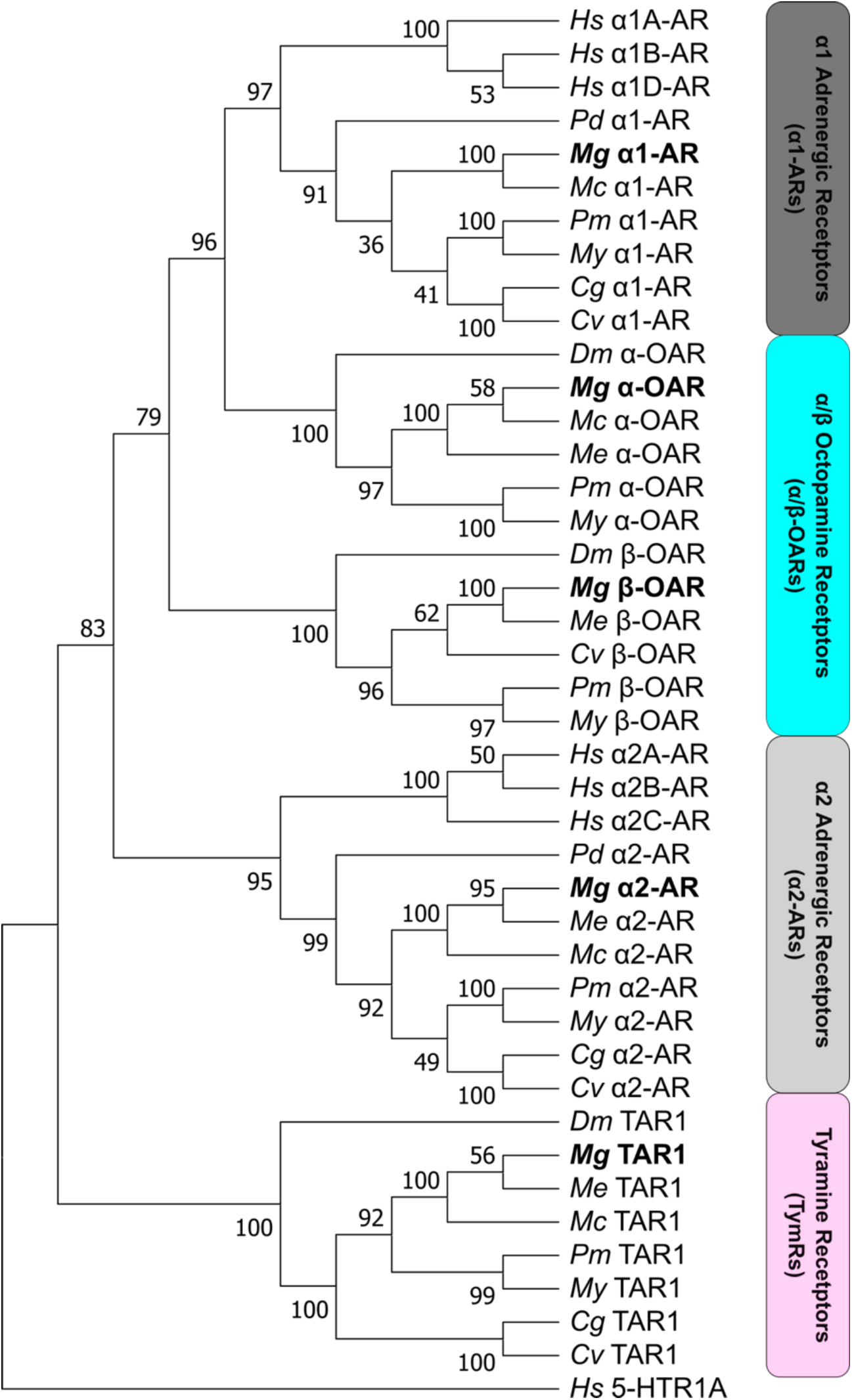
Orthology sequence analysis of adrenergic, octopaminergic and tyraminergic receptors proteins in *M. galloprovincialis* using human, fruit fly, and Dumeril’s clam worm (*Platynereis dumerilii*) sequences as references. Human 5-HTR1A was used as outgroup. Clades are coloured as follows: α1-Ars in dark grey, OARs in blue, α2-Ars in light grey, and TARs in pink. *M. galloprovincialis* sequence is highlighted in bold. Other bivalve species were included as previously described in the methods section. Abbreviations of species and enzymes are listed in Supplementary Table 10. Tree of adrenergic, octopaminergic and tyraminergic receptors. The tree was realised by using Maximum Likehood method and Le Gascuel model; bootstraps values were inferred from 1000 replicates. Initial tree(s) for the heuristic search were obtained automatically by applying Neighbor-Join and BioNJ algorithms to a matrix of pairwise distances estimated using the JTT model, and then selecting the topology with superior log likelihood value. A discrete Gamma distribution was used to model evolutionary rate differences among sites (5 categories (+G, parameter = 2.3316)). This analysis involved 42 amino acid sequences. All positions with less than 85% site coverage were eliminated.

**Supplementary Figure 9.**
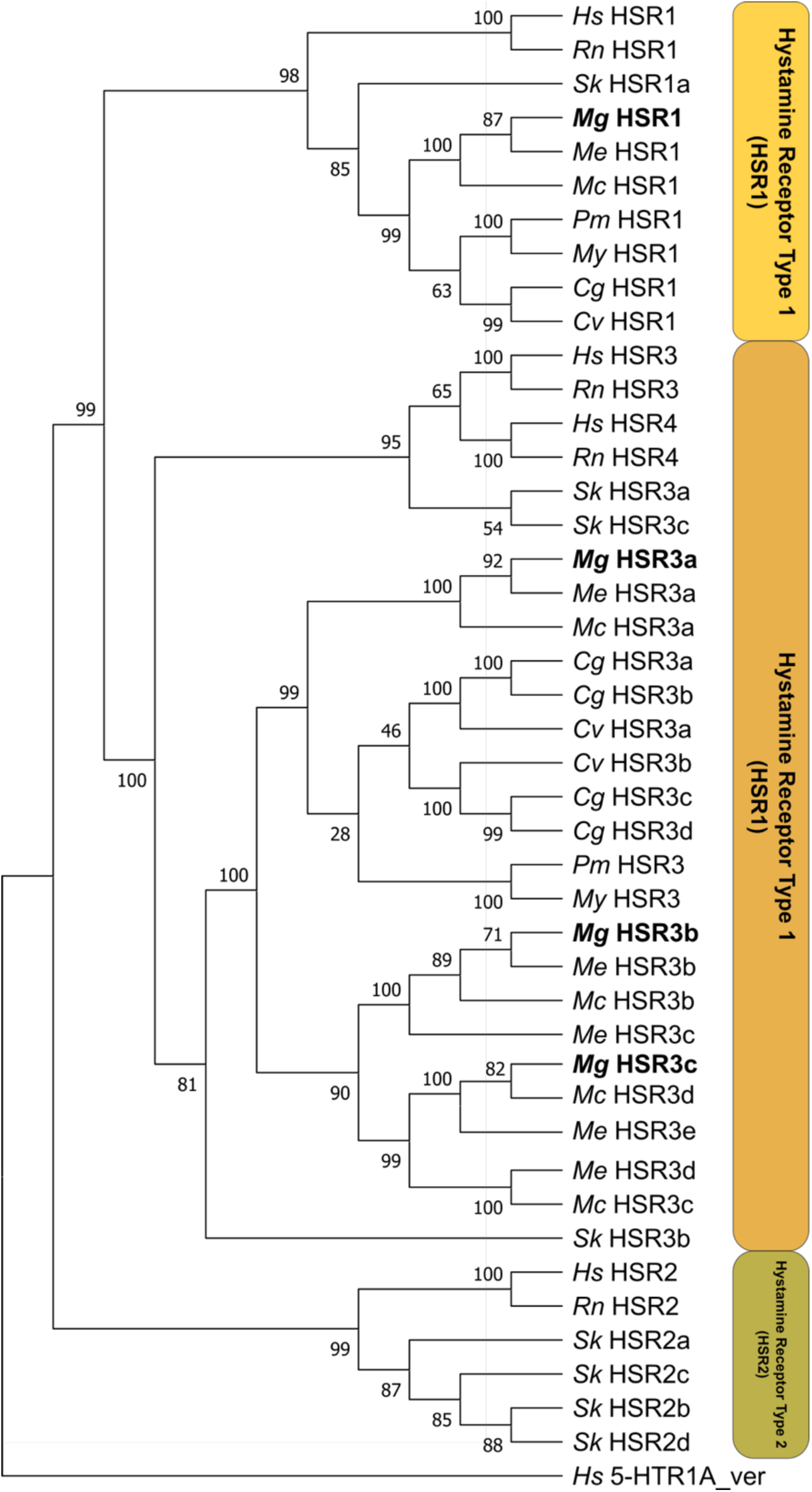
Orthology sequence analysis of HSRs proteins in *M. galloprovincialis* using human, rat, and acorn worm (*Saccoglossus kowalevskii*) sequences as references. Human 5-HTR1A was used as outgroup. HSRs subtypes are coloured with different shades of green. *M. galloprovincialis* sequence is highlighted in bold. Other bivalve species were included as previously described in the methods section. Abbreviations of species and enzymes are listed in Supplementary Table 11. The tree was realised by using Maximum Likehood method and Le Gascuel model; bootstraps values were inferred from 1000 replicates. Initial tree(s) for the heuristic search were obtained automatically by applying Neighbor-Join and BioNJ algorithms to a matrix of pairwise distances estimated using the JTT model, and then selecting the topology with superior log likelihood value. A discrete Gamma distribution was used to model evolutionary rate differences among sites (5 categories (+G, parameter = 3.2968)). The rate variation model allowed for some sites to be evolutionarily invariable ([+I], 2.24% sites). The tree is drawn to scale, with branch lengths measured in the number of substitutions per site. This analysis involved 44 amino acid sequences. All positions with less than 85% site coverage were eliminated. HSRs different types are highlighted by different shades of orange.

## Supplementary Tables

**Supplementary Table 1**. Table of HCR probes utilised to localise monoaminergic genes., The name of gene is reported in the first column, the gene ID is reported in the second column, andcolum three and four report, the amplifier used and the probes of the genes, respectively.

**Supplementary Table 2.** Table of HCR probes used to localise tissue marker genes. Table structure is the same of Supplemeantary Table 1.

**Supplementary Table 3.** List of sequences used for orthology assessment of decarboxylases proteins including AADC, HDC, and TDC. Human and fruit fly reference sequences (62) were used to retrieve AADC, HDC, and TDC in *M. galloprovincialis*, and human glutamate decarboxylases (GADs) were used to root the tree. Other bivalve species were included as previously described in the methods section. Asterisks in the table indicate presence in the protein of an additional domain.

**Supplementary Table 4.** List of sequences used for orthology assessment of copper containing hydroxylase. Humans, and murine DβH, human MODX, fruit fly TβH (14), great pond snail (*Lymnaea stagnalis*), whiteleg shrimp (*Litopenaeus vannamei*), and *Azumapecten farreri* DβH (101, 102) sequences were used to retrieve DβH and DβH-like in *M. galloprovincialis.* Human peptidylglycine alpha-hydroxylating monooxygenase (PAM) was used as outgroup. Other bivalve species were included as previously described in the methods section.

**Supplementary Table 5.** List of sequences used for orthology assessment of (MAOs. Human and murine MAO sequences were used to retrieve MAO in *M. galloprovincialis*. Human and murine Spermine Oxidase (SMOX) were used as outgroup to root the tree. Other bivalve species were included as previously described in the methods section.

**Supplementary Table 6.** Classification of MAOs based on presence or absence of conserved amino acids in their sequences. In case of similar amino acid, sequences were named *MAO-A- or MAO-B-like*. Sequences were named *MAO-like* in absence of conserved amino acids.

**Supplementary Table 7.** List of sequences used for orthlogy assessment of VMATs. Humans and fruit fly VMAT sequences were used to retrieve VMAT in *M. galloprovincialis*. Fruit fly and Western honey bee (*Apis mellifera*) orphan vesicular neurotransmitter transporter (portabella) were used as outgroup (65). Other bivalve species were included as previously described in the methods section.

**Supplementary Table 8.** List of sequences used for orthology assessment of 5-HTRs. Human, and fruit fly (23) sequences were used to retrieve 5-HTRs in *M.* galloprovincialis. Human rhodopsin, and fruit fly FMRFamide receptors were used to root the tree. Sequences presenting a nonspecific hit domain were called 5-HTRx-*like*. Other bivalve species were included as previously described in the methods section.

**Supplementary Table 9.** List of sequences used for orthlogy assessment of DRs. Human, and fruit fly sequences (66) were used to retrieve DRs in *M. galloprovincialis*. Fruit fly GluR was used to root the tree. Other bivalve species were included as previously described in the methods section. Sequences presenting a nonspecific hit domain were called DRx-like.Asterisks in the table indicate the absence of the specific domain (i.e.: 7tm_GPCRs super family) or the presence in the protein of an additional domain.

**Supplementary Table 10.** List of sequences used for orthology assessment of ARs, TARs, and OARs. Human, fruit fly, and Dumeril’s clam worm (27) were used to retrieve the previously mentioned receptors in *M. galloprovincialis*. Human 5-HTR1A was used to root the tree. Other bivalve species were included as previously described in the methods section.

**Supplementary Table 11.** List of sequences used for orthology assessmentHSRs. Human, rat, and acorn worm (*Saccoglossus kowalevskii*) (67) were used to retrieve HSRs in *M. galloprovincialis*. Human 5-HTR1A was used to root the tree. Other bivalve species were included as previously described in the methods section.

